# An accumulation-of-evidence task using visual pulses for mice navigating in virtual reality

**DOI:** 10.1101/232702

**Authors:** Lucas Pinto, Sue Ann Koay, Ben Engelhard, Alice M. Yoon, Ben Deverett, Stephan Y. Thiberge, Ilana B. Witten, David W. Tank, Carlos D. Brody

## Abstract

The gradual accumulation of sensory evidence is a crucial component of perceptual decision making, but its neural mechanisms are still poorly understood. Given the wide availability of genetic and optical tools for mice, they can be useful model organisms for the study of these phenomena; however, behavioral tools are largely lacking. Here, we describe a new evidence-accumulation task for head-fixed mice navigating in a virtual reality environment. As they navigate down the stem of a virtual T-maze, they see brief pulses of visual evidence on either side, and retrieve a reward on the arm with the highest number of pulses. The pulses occur randomly with Poisson statistics, yielding a diverse yet well-controlled stimulus set, making the data conducive to a variety of computational approaches. A large number of mice of different genotypes were able to learn and consistently perform the task, at levels similar to rats in analogous tasks. They are sensitive to side differences of a single pulse, and their memory of the cues is stable over time. Moreover, using non-parametric as well as modeling approaches, we show that the mice indeed accumulate evidence: they use multiple pulses of evidence from throughout the cue region of the maze to make their decision, albeit with a small overweighting of earlier cues, and their performance is affected by the magnitude but not the duration of evidence. Additionally, analysis of the mice's running patterns revealed that trajectories are fairly stereotyped yet modulated by the amount of sensory evidence, suggesting that the navigational component of this task may provide a continuous readout correlated to the underlying cognitive variables. Our task, which can be readily integrated with state-of-the-art techniques, is thus a valuable tool to study the circuit mechanisms and dynamics underlying perceptual decision making, particularly under more complex behavioral contexts.

## INTRODUCTION

Making decisions based on noisy or ambiguous sensory evidence is a task animals must face on a daily basis. Take, for instance, a mouse in the wild, whose navigation behavior relies on vision (Alyan and Jander, 1994/8; Etienne et al., 1996; Stopka and Macdonald, 2003). Amidst tall grass, deciding a route to a partially occluded food source (say, a corn plant) might require gradual accumulation of visual evidence, i.e. short glimpses of what may or may not be part of that plant. This example also highlights another important point about decision-making, namely that it is often performed in conjunction with other complex behaviors and can itself be a dynamic process occurring over seconds-long timescales. Here, the mouse must find its food source while navigating in a potentially changing environment; the corn plant may turn out to be a scarecrow, and evidence for or against this is typically used to interactively update a motor plan.

How the brain gradually accumulates sensory evidence has been the topic of extensive studies performed primarily in primates (Gold and Shadlen, 2007). However, much remains unknown regarding which brain areas are involved, and the specific circuit mechanisms and dynamics underlying this computation (Brody and Hanks, 2016). More recently, several groups have started using rodents to tackle such questions (Brunton et al., 2013; Carandini and Churchland, 2013; Hanks et al., 2015; Licata et al., 2017; Morcos and Harvey, 2016; Odoemene et al., 2017; Raposo et al., 2014; Scott et al., 2015). Rodents provide many complementary advantages to the use of primates, such as lower cost, larger scalability, and, particularly for mice, the wide availability of an ever-expanding arsenal of tools to record from and manipulate circuits with great spatiotemporal and genetic specificity in behaving animals (Chen et al., 2013; Deisseroth, 2011; Dombeck et al., 2007; Guo et al., 2014; Luo et al., 2008; Rickgauer et al., 2014; Sofroniew et al., 2016; Song et al., 2017; Svoboda and Yasuda, 2006).

Motivated by the above, we have developed a novel behavioral task in which head-fixed mice are required to gradually accumulate visual evidence as they navigate in a virtual T-maze. The side on which the majority of the evidence appears informs them of which of the two arms the reward is located in. Compared to freely moving behaviors, the use of virtual reality (VR) (Harvey et al., 2009) allows for better control of sensory stimuli, ease of readout of motor output, and, crucially, the head fixation required for many state-of-the-art optical techniques (Dombeck and Reiser, 2012; Minderer et al., 2016). In studying perceptual decision-making in conjunction with navigation, we emulate a more naturalistic context of rodent behavior. As brains are highly nonlinear systems that may engage qualitatively different mechanisms in different contexts, trying to approximate such conditions is arguably an important component towards understanding neural codes (Carew, 2005; Krakauer et al., 2017). Another highlight of our task is the use of multiple short pulses of sensory stimuli that are randomly distributed per trial according to Poisson statistics (Brunton et al., 2013; Scott et al., 2015). The diverse yet well-controlled nature of this stimulus set allows for the use of powerful computational approaches when analyzing the data (Brunton et al., 2013; Erlich et al., 2015; Hanks et al., 2015; Scott et al., 2015). Specifically, the stimuli are designed to be delivered in perceptually distinct pulses (“cues”), enabling neural recording and perturbation studies to trace/modulate precisely timed inputs into the animal’s brain. The randomized locations of the cues decorrelates the dynamics of evidence streams from the general progression of time, on a trial-by-trial basis, allowing us to investigate the distinct contributions of the amount and the timing of incoming evidence. This, in turn, gives us a better handle on the behavioral strategies the animals employ.

Here we perform a thorough characterization of various performance indicators, behavioral strategies and navigational aspects of the task, with the goal of providing a bedrock for future studies investigating the neural mechanisms underlying this behavior. We show that mice can consistently learn this task and solve it by using multiple pulses of visual cues distributed throughout the cue presentation period, thus accumulating evidence towards a decision. Moreover, we show that their performance is influenced by the magnitude of the evidence but not its duration. We also describe an intriguing, if small, tendency to alternate choices after rewards, and present logistic regression models that combine evidence and trial history as tools to quantify the behavior. Finally, we capitalize on the navigational component of the task and show that trajectories, though fairly stereotyped, may provide an ongoing readout correlated with cognitive variables.

## MATERIALS AND METHODS

### Animals and surgery

All procedures were approved by the Institutional Animal Care and Use Committee at Princeton University. Experiments were performed on both male and female mice aged 2 – 12 months, from several strains:

- 5 wild types [C57BL6/J, Jackson Laboratories, stock # 000664]
- 14 VGAT-ChR2-EYFP [B6.Cg-Tg(Slc32a1-COP4*H134R/EYFP)8Gfng/J, Jackson Laboratories, stock # 014548] (Zhao et al., 2011).
- 16 triple transgenic crosses expressing GCaMP6f under the CaMKIIα promoter, from the following two lines: Ai93-D;CaMKIIα-tTA [Igs^7tm93.1(tetO-GCaMP6f^)Hze Tg(Camk2a-tTA)1Mmay/J, Jackson Laboratories, stock # 024108] (Madisen et al., 2015); Emx1-IRES-Cre [B6.129S2-Emx1^tm1(cre)Krj/^J, Jackson Laboratories, stock # 005628] (Gorski et al., 2002).
- 8 Thy1-GCaMP6f [C57BL/6J-Tg(Thy1-GCaMP6f)GP5.3Dkim/J, Jackson Laboratories, stock # 028280] (Dana et al., 2014).
- 1 Thy1-YFP-H [B6.Cg-Tg(Thy1-YFP)HJrs/J, Jackson Laboratories, stock # 003782] (Feng et al., 2000).
- 6 DAT-IRES-CRE [B6.SJL-*^Slc6a3tm1.1(cre)Bkmn^*/J, Jackson Laboratories, stock # 006660] (Bäckman et al., 2006).

The various strains were part of different ongoing, unpublished studies, and are hereby grouped for behavioral analysis. Despite happening for technical reasons, the inclusion of and comparisons between different strains also allowed us to confirm that we can obtain comparable levels of behavioral performance across separate experiments and different experimenters. The mice underwent sterile stereotaxic surgery to implant a custom lightweight titanium headplate (~1 g, CAD design files available at https://github.com/sakoay/AccumTowersTools.git) under isoflurane anesthesia (2.5% for induction, 1.5% for maintenance). Briefly, after asepsis the skull was exposed and the periosteum removed using a bonn micro probe (Fine Science Tools) or sterile cotton swabs. The headplate was then positioned over the skull and affixed to it using metabond cement (Parkell). Some of the animals underwent additional procedures to either implant an imaging cranial window or make the skull optically transparent, as previously described (Guo et al., 2014; Harvey et al., 2012). Additionally, in the DAT-cre mice only, AAV5-EF1a-DIO-hChR2 (Penn Vector Core) was injected bilaterally in the ventral tegmental area (VTA) following standard biosafety level 1 procedures, and 300-µm optical fibers (Thorlabs) were implanted bilaterally above the VTA. The virus was injected as part of a separate study. The animals received one pre-operative dose of meloxicam for analgesia (1 mg/kg I.P. or S.C.) and another one 24h later, as well as peri-operative body-temperature I.P. saline injections to maintain hydration. Body temperature was maintained constant using a homeothermic control system (Harvard Apparatus). For cranial window implantation surgeries only, an intraperitoneal injection of dexamethasone (2-5 mg/kg) was given at the beginning of the procedure in order to reduce brain swelling. The mice were allowed to recover for at least 3 days before starting water restriction for behavioral training. They were then restricted to an allotted water volume of 1 – 2 mL per day, always ensuring that no clinical signs of dehydration were present and body mass was at least 80% of the initial value. If any of these conditions were not met, the mice received supplemental water (or had *ad libitum* access to water if more than mildly dehydrated) until recovering. Most typically, animals received their whole allotment during behavioral training, but received supplemental water if necessary, at least one hour after the end of training.

The animals were handled daily from the start of water restriction until they no longer showed any signs of distress, such as attempting to escape, defecating or urinating, which typically took 3 – 5 days. Mice were never picked up from the cage by their tails, instead voluntarily climbing onto the experimenter's hand or being gently lifted with a hand scooping movement. They were allowed to socialize in an enclosed enriched environment (~0.3 m2, with 5–10 mice) outside of behavioral sessions and before being returned to the vivarium at the end of each day.

### Behavioral task

#### Apparatus

We trained mice on virtual reality (VR) systems similar to ones described previously (Harvey et al., 2012; Low et al., 2014)(**Figure 1A**). Subjects were head-fixed using custom-made headplate holders and stood on a spherical treadmill comprised of a Styrofoam^®^ ball (8-inch diameter, Smoothfoam) placed on a custom 3D-printed cup and suspended by compressed air (60 – 65 p.s.i.). Compressed air was delivered through a 1.5 inch-diameter flexible hose (McMaster-Carr) coupled to an enclosed chamber beneath the cup. The source of air to this hose was first passed through a laminar flow nozzle (series 600 Whisperblast, Lechler), which dramatically reduced ambient noise by reducing air turbulence. The animals were placed on the ball such that their snouts were roughly aligned with the center of its upper surface, and at a height such that they could touch the ball with their whole forepaw pads, while not displaying noticeable hunching. This allowed them to run comfortably, with similar posture to when they are freely moving. A custom alignment tool that was mounted on the posts supporting the headplate holders was used to verify the mice's alignment with respect to VR system, and was critical to prevent side biases stemming from lateral asymmetries in controlling the ball (a CAD file for 3D printing the tool is available at https://github.com/sakoay/AccumTowersTools.git).

Ball movements controlled the mice's position within the VR environment, projected onto a custom-built Styrofoam^®^ toroidal screen with a 270˚ horizontal field of view, using a DLP projector (Optoma HD 141X) with a refresh rate of 120 Hz, a pixel resolution of 1024 × 768, and relative color balance of 0, 0.4 and 0.5, for the red, green and blue channels, respectively. Motion was detected by an optical flow sensor (ADNS-3080 Optical Flow Sensor APM2.6), coupled to infrared LED (890 nm, Digikey), and lying underneath the ball, within the cup on which the ball sat, which contained a 30-mm aperture covered with Gorilla Glass^®^ (Edmond Optics). Optical flow was transformed into displacement and output to the behavior control PC using custom code running on an Arduino Due (code and documentation may be downloaded from https://github.com/sakoay/AccumTowersTools.git). The accuracy of this measurement depends on the presence of sufficiently high-contrast features on the ball surface. In order to ensure this, the styrofoam balls were either roughened with steel wool or small black marks were made crisscrossing the entire area using a permanent marker. Treadmill displacements in *X* and *Y* (d*X*, d*Y*) resulted in equal translational displacements in the VR environment (i.e. gain of 1). To set the virtual viewing angle θ, the acute angle between the line formed by the displacement vector and the *Y* axis line was calculated as

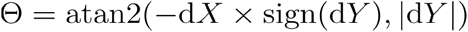

The rate of change in θ (radians/second) was then calculated using an exponential gain function of Θ, as follows:

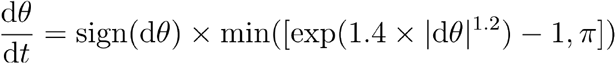

This gain was tuned to damp small values of dθ/dt, stabilizing trajectories in the maze stem where mice typically made only small course corrections to maintain forward movement. The exponential dependence ensured that mice could still perform sharp turns (i.e. generate large values of dθ/dt) into the maze arms; see **Supplementary Methods** for more details.

Reward delivery was controlled by TTL pulses from the control PC sent to a solenoid valve (NResearch) and done through a beveled plastic 100-µL pipette tip coupled to PVC plastic tubing (McMaster-Carr). Sounds were played through conventional computer speakers (Logitech). The apparatus was enclosed in a custom-designed cabinet (8020.inc) lined with sound-absorbing foam sheeting (McMaster-Carr). The whole system was controlled by a PC running the matlab-based software ViRMEn (Aronov and Tank, 2014) (available for download at https://pni.princeton.edu/pni-software-tools/virmen-virtual-reality-matlab-engine).

#### Accumulating-towers task

Mice were trained to run down a virtual T-maze (total length: 330 cm, visual width: 10 cm, allowed travel width: 1 cm, wall height: 5 cm) and retrieve a fluid reward from one of the two end arms (each measuring 10.5 × 11 × 5 cm, length × width × height) (**Figure 1B**). As they ran down the central stem they saw briefly-appearing, tall, high-contrast objects (towers, width: 2 cm, height: 6 cm) on either side of the maze, and the arm on the side with the most towers contained the reward. Towers appeared whenever the animals were 10 cm away from them, and disappeared 200 ms later. In each trial, tower position within the cue period (200 cm) was drawn randomly from spatial Poisson processes with means of 7.7 towers/m for the rewarded side and 2.3 towers/m for the non-rewarded (minority cue) side (i.e. an overall tower density of 5 m-1), and a refractory period of 12 cm; see **Supplementary Methods**.

At the start of each trial the mice were teleported to a 30-cm long starting location and the maze appeared. The virtual view angle was restricted to be 0 throughout this region, in essence acting as a buffer zone during which mice could straighten out their running patterns. After they ran past the starting location, the floor and wallpapers changed to indicate they were in the main part of the maze, and mice were then free to rotate the view angle. Towers could appear anywhere within the first 200 cm of the maze (cue period), and the last 100 cm of the maze (delay period) did not contain any towers but had the same wallpaper as the cue period. The wallpaper changed in the arms of the maze but was identical on both sides. After the mice reached one of the arms, the world was frozen for 1 s and then disappeared for 2 s (i.e. screen became black). A correct response was thus followed by a 3-s inter-trial interval and was rewarded with a drop of 10% (v/v) sweet condensed milk solution (4 – 8 µL), whereas an error was followed by a sound and an additional 9-s timeout period (total inter-trial interval of 12 s). Trials timed out after 600 sec (or 60 sec in some sessions).

Every session started with warm-up trials of a visually-guided maze. In this maze, towers appeared exclusively on one side, and a tall visual guide (30 cm) positioned in one of the arms indicated the reward location. In order to advance to the main maze, the mice were required to perform at least 10 warm-up trials at a minimum of 85% correct, with a maximum side bias (difference in percent correct between right- and left-rewarded trials) of 10% and at least 75% of good-quality trials, defined as trials in which the total distance traveled is at most 110% of the maze length. Once in the main maze, performance was constantly evaluated over a 40-trial running window, with two purposes. First, if performance fell below 55% correct, animals were automatically transferred to a block of trials in an easier maze, with towers shown only on the rewarded side, but with no visual guide. This block had a fixed length of 10 trials, after which the mouse returned to the main maze regardless of performance. The other purpose of the 40-trial window was to assess and attempt to correct side bias. This was achieved by changing the underlying probability of drawing a left or a right trial according to a balanced method described in detail elsewhere (Hu et al., 2009). In brief, the probability of drawing a right trial, *p_R_,* is given by

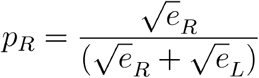

Where *e_R_* (*e_L_*) is the weighted average of the fraction of errors the mouse has made in the past 40 right (left) trials. The weighting for this average is given by a half-Gaussian with σ = 20 trials in the past, which ensures that most recent trials have larger weight on the debiasing algorithm. To discourage the generation of sequences of all-right (or all-left) trials, we capped 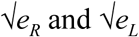 to be within the range [0.15,0.85]. In addition, a pseudo-random drawing prescription was applied to ensure that the empirical fraction of right trials as calculated using a σ = 60 trials half-Gaussian weighting window is as close to as possible, i.e. more so than obtained by a purely random strategy. Specifically, if this empirical fraction is above, right trials are drawn with probability 0.5 *p_R_*, whereas if this fraction is below, right trials are drawn with probability 0.5 (1 + *p_R_*).

The six DAT-IRES-Cre mice included in the dataset ran a slightly different version of the task. For these animals, the cue region was 220 cm and the delay was 80-cm long (vs. 200 and 100), the tower density was 3.5 m-1 (vs. 5), and the tower refractory period was 14 cm (vs. 12). For this reason, these mice were not included in any of the analyses except the comparison of performance between different strains (**Supplementary Figure 4**).

#### Shaping

Details about the shaping procedure can be found in **Supplementary Figure 1** and **Supplementary Table 1**. Briefly, mice underwent at least 11 shaping stages (T1 – T11, where T11 is the final maze explained in the previous paragraph). The first 4 stages (T1 – T4), consisted of visually-guided mazes with cues throughout the stem, and with progressively increasing lengths. Moreover, while the appearance of towers was triggered by proximity as previously explained, they did not disappear after 200 ms. Final length was reached at maze T4. Next, the visual guide was removed (T5) and the cue period length was progressively decreased to its final value of 200 cm (T6 – T7). Up to T7, towers always appeared only on the rewarded side. The next step in shaping was to progressively increase the rate of minority cues (i.e. towers on the non-rewarded side, T8 – T11) and finally to make the towers disappear after 200 ms (T10 – T11). An earlier version of the shaping procedure had 14 instead of 11 steps, whose only difference was to introduce changes more gradually, but eventually reaching an identical maze. These animals were included in all the analyses except that in **Supplementary Figure 1**. Mice were trained 5 – 7 days / week, for one 1-hour session per day. The only exceptions to this were for the first two days of training, where mice were acclimated to the VR setup for 30 and 45 minutes, respectively. Mice typically took 6 – 7 weeks to reach the final stage (see **Results**).

### Data Analysis

#### Data selection

The initial dataset was comprised of 1,067 behavioral sessions from 38 mice, with a total of 194,766 trials from the final accumulation maze (182.5 ± 2.2 trials/session, mean ± SEM). Besides regular behavioral training, we also included sessions occurring during either two-photon or widefield Ca2+ imaging, or optogenetic manipulation experiments. In the latter case, we only included control (laser off) trials (70 – 85% of trials in a session). Unless otherwise stated, we applied the following block-wise data inclusion criteria: 1) whole trial blocks (i.e. consecutive trials in the same maze, of which there could be multiple in a session) with an overall performance of at least 60% correct, including trials of all difficulties; 2) trials with a maximal traveled distance of 110% of nominal maze length (Harvey et al., 2012); 3) no timed-out or manually aborted trials; and 4) after applying criteria 1 – 3, individual mice with at least 1,000 trials. We thus selected 135,824 trials from 878 sessions and 25 mice (mean ± SEM: 5,433 ± 774 trials/mouse, range: 1,118 – 15,283; mean ± SEM: 35.1 ± 4.5 sessions/mouse, range: 7 – 86). For analyses involving effects of trial history (**Figures 7A–F**), we excluded all optogenetic sessions to avoid the use of non-consecutive trials, as well as those without at least 5 trials of history (i.e. first five trials of a block). Those additional criteria yielded 66,411 trials from 18 mice and 507 sessions. For all model fits except the SDT model (**Figures 4, 6, 7, –7, Supplementary Figures 5, 6**), we required one trial of history, to allow for fair comparison between models with and without trial history. Those criteria yielded 81,705 trials from 20 mice and 597 sessions.

#### Psychometric curves

We built psychometric curves by plotting the percentage of trials in which the mouse chose the right arm as a function of the difference in the number of right and left towers (#R – #L, or Δ). For **Figure 2A** and **Supplementary Figure 6C, D**, Δ was binned in groups of 3 and its value defined as the average Δ weighted by the number of trials. We fitted the psychometric curves using a 4-parameter sigmoid:

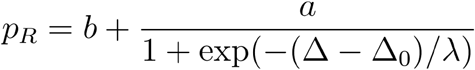

The slope of this sigmoid (**Figure 2C**) was defined as *a*/4λ (the derivative of the curve at Δ_0_), and lapse rate (**Figure 2D**) was defined as the average error rate (%) in all trials with |Δ| ≥ 10.

#### Logistic regression analysis

To assess how evenly mice weighted sensory evidence from different segments of the cue period (**Figures 3A, B**), we performed a logistic regression analysis in which the probability of a right choice was predicted from a logistic function of the weighted sum of the net amount of sensory evidence per segment, Δ(*y*), where *y* is one of 5 equally spaced segments between 10 and 200 cm (because tower appearance was triggered by proximity, the earliest possible tower occurred at *y* = 10 cm):

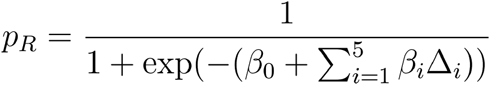

Note that this analysis is similar to the commonly used reverse correlation (e.g., (Brunton et al., 2013). We have confirmed that both analyses yield very similar results (not shown). To estimate the amount of recency or primacy effects from the logistic regression coefficients (**Figure 3C**) we computed a weight decay ratio as [(Δ_4_ + Δ_5_)/2] / [(Δ_1_ + Δ_2_)/2], such that values smaller than 1 indicate primacy effects (i.e. initial portions of the cue period are weighted more towards the decision) and 1 indicates spatially homogeneous accumulation. To calculate the significance of the decay ratio for each mouse, spatial bin identities for each trial were shuffled 200 times, and in each iteration the logistic regression model was refit, yielding a null distribution for the ratio. *P*-values were calculated as the proportion of shuffling iterations whose decay ratio was smaller than the actual ratio. Errors on logistic regression coefficient estimates for individual mice were calculated as the standard deviation of the distribution given by sampling the trials with replacement and refitting the model 200 times.

#### Effect of number of towers, cue and delay duration

For this analysis, |Δ| and total number of towers were binned into groups of two, and effective duration of cue and delay periods into 10-cm bins. Effective cue period duration was defined as the difference in the position of the last and first tower, regardless of side, and effective delay duration was defined as 300 (stem length in cm) minus the position of the last tower. We first calculated performance (% correct) separately for each binned value of |Δ|, as a function of either cue duration (**Figure 5A**), total number of towers (**Figure 5B**) or delay duration (**Figure 5D**). To better estimate the relative contributions of |Δ|, total number of towers and period duration (**Figure 5C**), we fit a linear model to the data as follows. First, for each mouse, we calculated performance for all 3-way combinations of binned predictor values (where period duration is of either cue or delay), and subtracted the average performance for that mouse. We then averaged these mean-subtracted performance values across mice, and fit a 3-parameter linear regression. Fitted parameter significance values were derived from the *t*-statistic of the parameter, i.e. its average divided by its standard deviation, which follows a *t* distribution with *n* – *p* – 1 degrees of freedom, where *n* is the number of data points and *p* is the number of parameters (Chatterjee and Hadi, 2015).

#### Trial history analysis

Alternation bias for each mouse (**Figures 67C–E**) was calculated as the percentage of trials in which they chose the arm opposite to their previous choice, subtracting the overall average percentage. In other words, we calculated the average difference between red and black, and blue and black curves in **Figures 7A, B**, with appropriate sign conventions. Note that positive alternation bias values indicate visiting the opposite arm in the following trial, whereas negative values indicate perseveration, i.e. visiting the same arm. For the analyses going five trials back (**Figures 7D, E**), bias is always defined with respect to trial zero (t_0_).

#### Analysis of running speed

Speed in cm/s was calculated on a trial-by-trial basis using the total x-y displacement for 0 < y < 300 cm (i.e. for the central stem). For the analysis in **Supplementary Figure 7C**, for each mouse, within-session standard deviation is the standard deviation across trials in the same session, averaged across sessions, and across-session standard deviation is the standard deviation of the distribution of average speeds for each session.

#### View angle analysis

In a given trial, the mouse traverses the T-maze with a y position trajectory y(t) that is not necessarily monotonically increasing, as variations in motor control can cause small amounts of backtracking. We therefore defined the view angle at a particular Y position, θ(Y), as the value of θ at the first time t at which y(t) ≥ Y. For the choice decoding analysis in **Figure 8B**, we defined an optimal choice decoding boundary θ_cd_(y) for a given y position by requiring that the fraction of right-choice trials with θ(y) > θ_cd_(y) be equal to the fraction of left-choice trials with θ(y) < θ_cd_(y). Thus, θ_cd_(y) is the boundary that most equally separates the right- vs. left-choice distributions. The choice decoding accuracy was defined as the percent of right-choice trials with with θ(y) > θ_cd_(y). For the analysis in **Figure 8D**, for each mouse we subtracted single-trial view angle trajectories from their average trajectory, separately for left and right choice trials. We then calculated tower-triggered trajectories separately for right and left towers, where y = 0 was defined as the position of the mouse when the tower appeared.

#### Brunton et al model

For the analyses in **Figures 6A–C** and **Supplementary Figure 5**, we fit the model described in detail in (Brunton et al., 2013)). It is part of the family of the widely used drift diffusion models (DDMs) (Gold and Shadlen, 2007; Ratcliff and Rouder, 1998), and models a latent decision variable *a,* whose amount of change per maze y position is given by

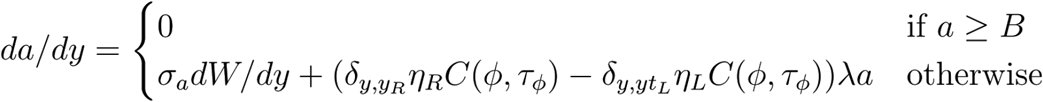

where *δ_y,yR_* and *δ_y,yL_* are delta functions at the spatial positions of right and left tower onset, *η* are i.i.d. variables drawn from *N*(1,*σ^2^_s_*), the initial value of *a* is drawn from *N*(1,*σ*^2^_*s*_), and *dW* is a *s a* Wiener process. *B* parametrizes the height of a sticky bound, *C* is a function of two parameters, *φ* and *τ_φ_*, and describes the adaptation dynamics to the sensory pulses. The memory time constant is given by *τ = 1/λ*. Finally, a bias parameter determines the position of the threshold above which a right choice is made and the lapse rate represents the probability of trials in which subjects will ignore the stimulus and choose randomly. Both these parameters are applied at the end of the trial when converting the continuous decision variable into a binary decision. The model was fit using a gradient descent algorithm to minimize the negative log likelihood cost function, using the interior-point algorithm from the Julia package Optim. Gradients were computed through automatic differentiation with respect to model parameters for each trial. Automatic differentiation makes it possible to efficiently compute complex derivatives with machine precision, and greatly improves the optimizer’s performance. We used the following parameter value constraints to fit the models: -5 < *λ* < 5, 0 < *σ*^2^_*s*_ < 200, 0 < *σ*^2^_*s*_ < 200, 0 < *σ*^2^_*i*_ < 30, 5 < *B* < 25, 0 < *φ* < 1.2, 0.001 < *τ_φ_* < 2 (maximum length of the cue period), -5 < bias < 5, 0 < lapse < 1.

#### Signal detection theory (SDT) model

We used the SDT model (**Figure 6D**) developed by (Scott et al., 2015)), where details about the method can be found. Briefly, the probability of making a correct choice (*pc*) in a given trial was modeled as the difference of two Gaussian distributions given by unique tower counts on the sides with the larger and smaller number of towers, *L* and *S*, where the variance of the distributions were the free parameters *σ*^2^_*L*_ and *σ*^2^_*L*_:

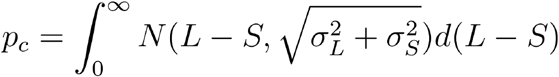

where *L* and *S* are an integer number of towers between 0 and 15 (we excluded trials with 16 or more towers on one side since there were very few of them). The model thus had 16 free parameters, whose best fit values were the ones that maximized the likelihood of the mice's choices using the Matlab function fmincon's interior-point algorithm. To fit this model, we only used the metamouse (aggregate) data, since individual mice had too few trials to obtain good fits.

In order to statistically distinguish between the linear variance and the scalar variability hypotheses, we explicitly modeled *σ* as either a linear or a square root function of the number of towers instead of fitting separate *σ* values. Specifically, we fit two separate two-parameter models to the data, one where *σ*(*n*) = *β*_0_ + *β_1_n* and another where *σ*^2^(*n*) = *β*_0_ + *β_1_n*, with *β*_0_ and *β*_1_ being free parameters and requiring *β*_0_ ≥ 0 (similar to models b and d in Fig. 4 of Scott et al., 2015). Statistical significance was calculated by bootstrapping the data 1000 times and defined as the proportion of bootstrap experiments in which the linear variance model had better goodness-of-fit than the scalar variability model (using the model information index, see below). For the two-parameter models, we also fitted individual mouse data. Note that for all SDT models, unlike the other models, we used the full dataset including non-contiguous trials (n = 25 mice, 135,824 trials), in order to gain statistical power.

#### Heuristic models with trial history

We fitted logistic regression models (**Figures 7G, H** and **Supplementary Figure 6**) where the probability of making a right choice, *p_R_*, was a function of both sensory evidence (with two different parameterizations, see below), and trial history, as follows:

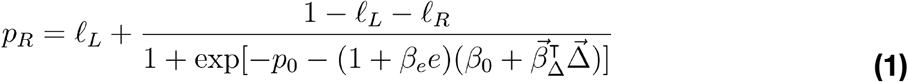

In the equation above, *p*_0_ ≐ −ln(1/*f*_R_ − 1) where *f_R_* is the fraction of right-choice trials in the given dataset. This was introduced such that when all the free model parameters 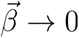, then *p_R_* → *f_R_*, i.e. the models considered here are a nested set w.r.t. the null hypothesis that the mouse has a constant right-choice probability *f_R_*, which facilitates model comparison. ℓ_*R*_(ℓ_*L*_) are lapse rates and can be interpreted as the probability of the mouse making a right choice in very easy right- (left-) rewarded trials, which can depend on both the mouse’s previous choice and the resulting outcome of the previous trial (**Figures 7A, B**). In other words, because of the observed trial-history-dependent vertical shifts in the psychometric curves (**Figures 7A, B, Supplementary Figures C, D**), history-dependent terms modulated the lapse rates and not the evidence terms inside the logistic function. Because probabilities must be bounded such that 0 ≤ *p_R_* ≤ 1, we constrained both lapse rates to be in the range 0 ≤ ℓ*R/L* ≤ 0.5 by applying a cosine transform to the otherwise linear model of dependence on history terms 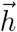:

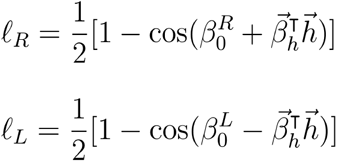

Here, 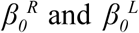 are history-independent lapse rates, and 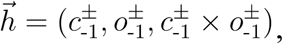 where the previous-choice indicator function 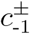 is defined to be +1 (-1) if the mouse chose right (left) in the previous trial, and the previous-outcome indicator function 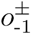 is defined to be +1 (-1) if the mouse was rewarded in the previous trial. Back to equation **(1)**, *β_e_* was introduced to account for history effects that change the slope of the evidence dependence after errors (**Figure 7B**), multiplying the “error” indicator function *e*, which is defined to be +1 if the mouse made a wrong choice in the previous trial and -1 otherwise. Finally, 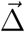 is a vector of evidence weights that took two different forms. For the model in **Figure 7H**, this vector was equivalent to that in the spatial bin logistic regression model described previously, i.e. the cue region was divided into 5 equal-width bins spanning y = 10 – 200 cm, and 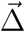 was set to be a 5-dimensional vector where each coordinate corresponds to #R – #L towers in each of the bins. For the model in **Supplementary Figure 6** (cue order model), we built the evidence vector as follows. For a given trial, cues (including both sides) were first ranked by their y position in ascending order, i.e. rank 1 corresponds to the first cue seen by the mouse on that trial. To improve statistical power, this rank was downsampled by a factor of 3 before defining 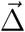 as a vector of *#R – #L* restricted to cues with the corresponding ranks. That is, the first coordinate of 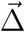 is *δ*_1-3_ = [*#R - #L*]_ranks 1-3_, the second coordinate is *δ*_4-6_ = [*#R - #L*]_ranks 4-6_, and so forth. Because the total number of cues differs from trial to trial due to random sampling, this means that not all trials have information for what would be the 2^nd^ and onwards coordinates of 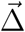. In order for the model to be well-posed, the dimensionality of 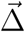 is fixed to be the maximum possible such that there are at least 50 trials that have information for the last coordinate. Trials with fewer than this number of cues are therefore assumed to have Δ_*i*_ = 0 for the remaining coordinates. The evidence vector was then normalized as 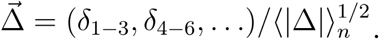. The normalization factor 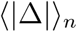 depends on the number of cues *n* for the given trial, and defined to be the average |*#R – #L*| over all trials in the dataset with the same number *n* of cues. These models were fit by maximizing the log-likelihood including L1 penalty terms for all free parameters (Schmidt, 2010). For a more detailed treatment of the models and fitting procedure, refer to **Supplementary Methods**.

#### Alternative strategy models

**1) Trial-history-only model (Figure 4A)**: we fitted a logistic regression model in which choice was a function only of previous trial history. The model is the same as in equation (1), except it did not contain the spatially binned evidence terms 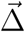. **2) *K* random tower models**: for these models, we assume that in each trial the mouse chooses *k* towers at random (from all presented towers) and uses only those to make its decision. The probability of making a right choice is thus equivalent to the probability that the majority of the *k* towers is on the right side, i.e. *kR* > *k*/2. This is given by the hypergeometric distribution, which gives us the probability of *k*/2 in *k* random draws without replacement from a population of size #R + #L, given that we know that #R towers are on the right. We implemented this using the Matlab function hygecdf. Note that in the special case where *k* = 1 (i.e. the one random tower model, **Figures 4B,C**), the probability of choosing right reduces to #R/(#R+#L). **3) First tower** and **Last tower** models: here we simply assume that the mouse will choose right if the first (last tower) appears on the right. In order to account for lapse rates in model classes **2)** and **3)**, the obtained probability of going right, *p_R_^m^*, was modulated by the experimentally measured lapse rates for right and left trials (*l_R_ and l_L_*) for each animal, i.e. 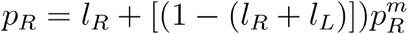. These lapse rates can be assessed independently of 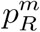 by using trials where there are only towers on one side, for which 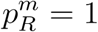 for all these models. Therefore, *l_R_*(*l_L_*) is the fraction of error trials out of all trials with only left (right) towers. For further details on alternative strategy models, refer to **Supplementary Methods**.

#### Model comparison and cross-validation

All models except the SDT were evaluated separately for each mouse using 70 runs of 3-fold cross-validation (i.e. using ⅔ of the data for training and ⅓; for testing the model), making sure that the subsamples of data used in each run were identical for all models. In these models, we are either hypothesizing that trial history effects do not exist, or that trial history effects have particular explicit parameterizations as described above. Therefore the model prediction for each trial is uncorrelated with that of any other trial (beyond any explicitly modeled history effects), and for each run we calculated the log likelihood (ln *L*) of the test dataset given the best-fit parameters on the training set, as follows. Let the mouse’s choice on the i^th^ trial be which *c_i_, i* = 1, …, *m* is 1 (0) if the mouse chose right (left). The likelihood of observing this choice is given by the binomial distribution 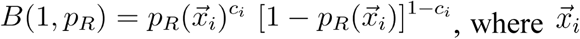 are various features of the ith trial that the model depends on (evidence, trial history, and so forth). Taking the product of individual-trial likelihoods we obtain:

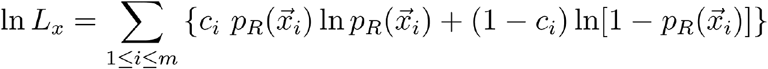

Additionally, we calculated a reference log likelihood (ln *L_0_*) of a trivial model with constant probability of going right, *f_R_*, being the experimentally-measured fraction of trials in which the animal went right. We then defined the goodness-of-fit of a given model to be the model information index, *MI*:

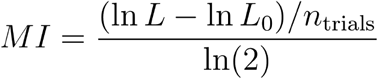

In other words, we calculated a trial-averaged excess likelihood of the model (compared to the trivial model) and converted the log to base 2, which gives us the amount of information of the model in units of bits/trial (Paninski et al., 2004; Pillow et al., 2008). In the cross-validation framework, all model parameters are extracted using the training set of 2/3rds of trials, and the *MI* values are evaluated only using the test set of 1/3rd of trials, which penalizes overfitting. To compare models across the population, we took the median *MI* across the 210 cross-validation runs for each mouse. To assess significance of the difference between *MI*s for a given mouse, we calculated a *P* value as the proportion of runs in which one model out(under)-performed the other.

#### General statistics

Datasets were tested for normality using the Lilliefors' modification of the Kolmogorov-Smirnov test. For comparisons between two normally distributed datasets, we used two-sided t-tests, and for non-normally distributed datasets we used the Wilcoxon sign rank test (or their paired test counterparts where appropriate). Multiple comparisons were corrected using the false discovery rate correction method described in (Benjamini and Hochberg, 1995) (see also (Guo et al., 2014)). Briefly, *P* values are ranked in ascending order, and the *i*th ranked *P-*value, *P_i_*, is deemed significant if it satisfies *Pi* ≤ (*αi*)/*n*, where *n* is the number of comparisons and *α* = 0.05 in our case. For tests involving the comparisons among multiple groups, we performed one- or two-way ANOVAs with repeated measures, followed by Tukey's post-hoc tests where appropriate. Binomial confidence intervals were calculated as 1-σ intervals using Jeffrey’s method.

## RESULTS

### Accumulating-towers task

We have developed a novel pulse-based evidence accumulation task for mice navigating in virtual reality (VR) (**Figure 1, Supplementary Movie 1**). Briefly, mice were trained to navigate on a virtual T-maze to retrieve water rewards from one of the two arms. While they ran down the central part of the maze (cue region, 200 cm), salient visual cues (towers) appeared transiently (200 ms) on either side. After a delay period without any cues (100 cm), the animal made either a right or left turn into one of the arms, and was rewarded if this corresponded to the side that had the highest number of towers (**Figure 1B**). Incorrect choices led to the playing of an error-indicating sound and a time-out period of 9s, in addition to the regular 3-s intertrial interval. The cues were distributed as spatial Poisson processes (**Supplementary Methods**) with different rates on the rewarded and unrewarded sides, such that the positions and number of towers on either side varied from trial to trial. This, together with the transient nature of the cues, meant that towers needed to be incrementally accumulated towards a decision. Importantly, the precisely controlled stimulus times allowed for powerful computational approaches when analyzing the data.

We developed detailed shaping procedures whereby different elements of the final task were gradually introduced, with well-defined and automated criteria for progression through the various stages (**Supplementary Figure 1, Supplementary Table 1**). Most animals in the dataset underwent an 11-step procedure, taking 34.8 ± 4.5 daily sessions (mean ± SEM, n = 17 mice) to reach the final stage, or 6 – 7 weeks including training breaks.

**Figure 1.**
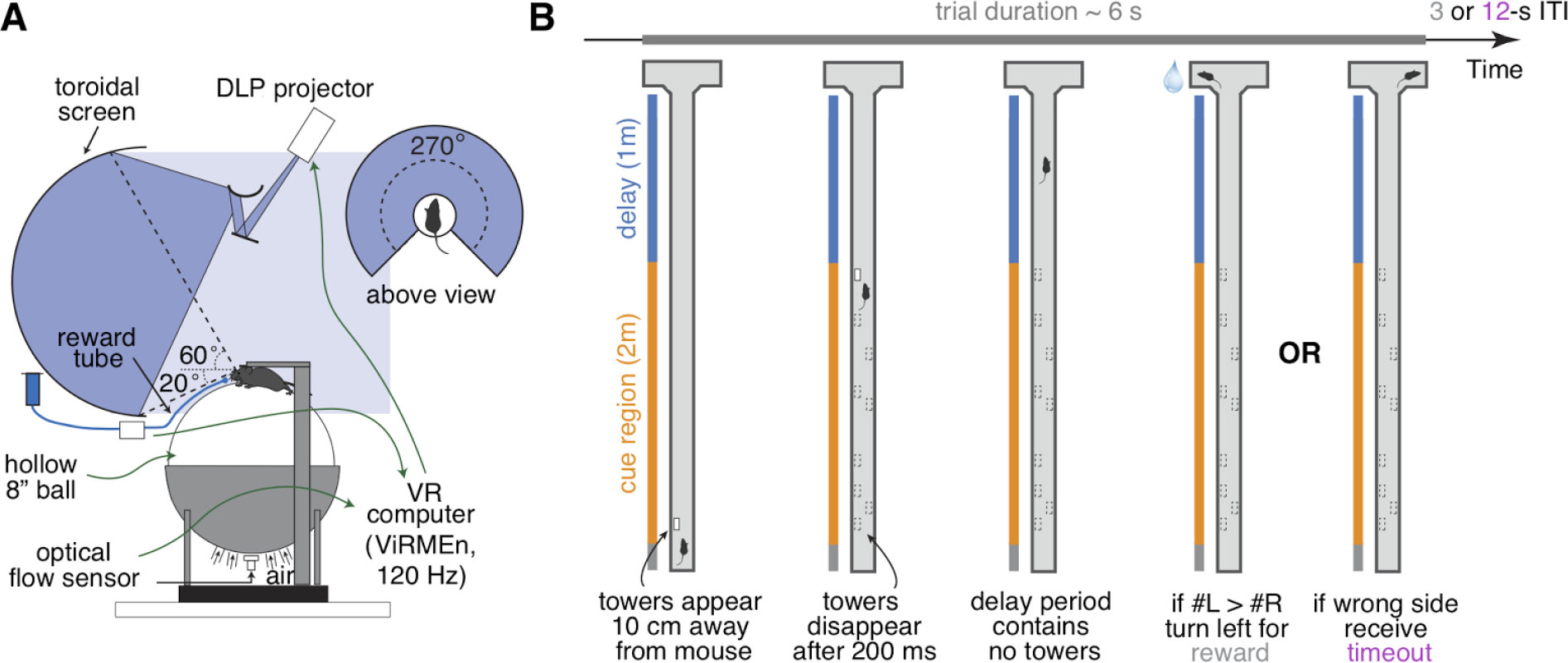
Accumulating-towers task. **(A)** Schematic drawing of the VR setup used to train mice in the task. **(B)** Schematic drawing of the task showing the progression of an example left-rewarded trial.

### Mice are sensitive to the amount of sensory evidence

We analyzed data from 25 mice with at least 1,000 trials on the final accumulation maze, obtaining a total of 135,824 trials (see **Materials and Methods** for details on data selection). Overall task performance including all trial difficulties was 68.7 ± 0.5 % correct (mean ± SEM, n = 25, range: 64.7 – 72.8 %). We were able to obtain many good-performance sessions for most mice (**Supplementary Figure 2**), including very high-performance sessions with steep psychometric curves and low lapse rates (e.g. **Figure 2A**). Performance was variable across sessions (standard deviation of overall performance across sessions: 7.0 ± 0.4 %, mean ± SEM across mice), and could fluctuate within sessions (**Supplementary Figure 2C** and **Supplementary Figure 3**), typically dropping towards the end, presumably when the mice were sated (**Supplementary Figure 3A**).

Importantly, performance was modulated by the amount of sensory evidence (#R – #L towers, or Δ), as revealed by psychometric curves (**Figures 2A, B, Supplementary Figure 3B**). Taking all blocks of consecutive trials in the same maze level into account (a block is defined here as consecutive trials in the same maze level, and there could be multiple within a session, see **Materials and Methods**), the slope of the psychometric function was 4.7 ± 0.2 %/tower (mean ± SEM, **Figure 2C**), and the lapse rate, defined as the error rate for |Δ| ≥ 10, was 21.4 ± 0.9 % (**Figure 2D**).

Given the variability we observed in performance, we next explored different performance selection criteria. For instance, if only the blocks over the 90th percentile of overall performance were selected, psychometric slope and lapse rate were 8.2 ± 0.4 %/tower and 12.8 ± 1.1 %, respectively (**Figures 2C, D**, green histograms). Of course, if we assume that different behavioral blocks are noisy samples from static psychometric curves, applying these criteria would trivially yield better performance indicators. To explicitly test for this possibility, we assumed that each mouse had a static psychometric curve across all sessions and generated 200 surrogate datasets by drawing samples from the binomial distributions given by the psychometric curves at the actual experienced values of Δ towers, and reselected the top 10% of blocks for each of these 200 draws. Interestingly, only the average improvements in lapse rate, but not psychometric slope, were significantly smaller in the surrogate data than the actual observed improvement (**Figures 2 E, F,** *P* = 2.4 × 10^−4^ and 0.97 respectively for lapse and slope, one-sided signed rank test; 7/18 mice have individually significant differences for lapse). This can be understood by noting that trials with |Δ| towers ≥ 10 comprise only ~10% of the total number, making them a relatively unimportant contribution to the overall performance and therefore not much affected by selection in the simulated static-psychometric data. We thus conclude that actual mice exhibit significantly lower lapse rates on high-performance behavioral blocks, but not more sensitivity to evidence, beyond that expected by random sampling.

**Figure 2.**
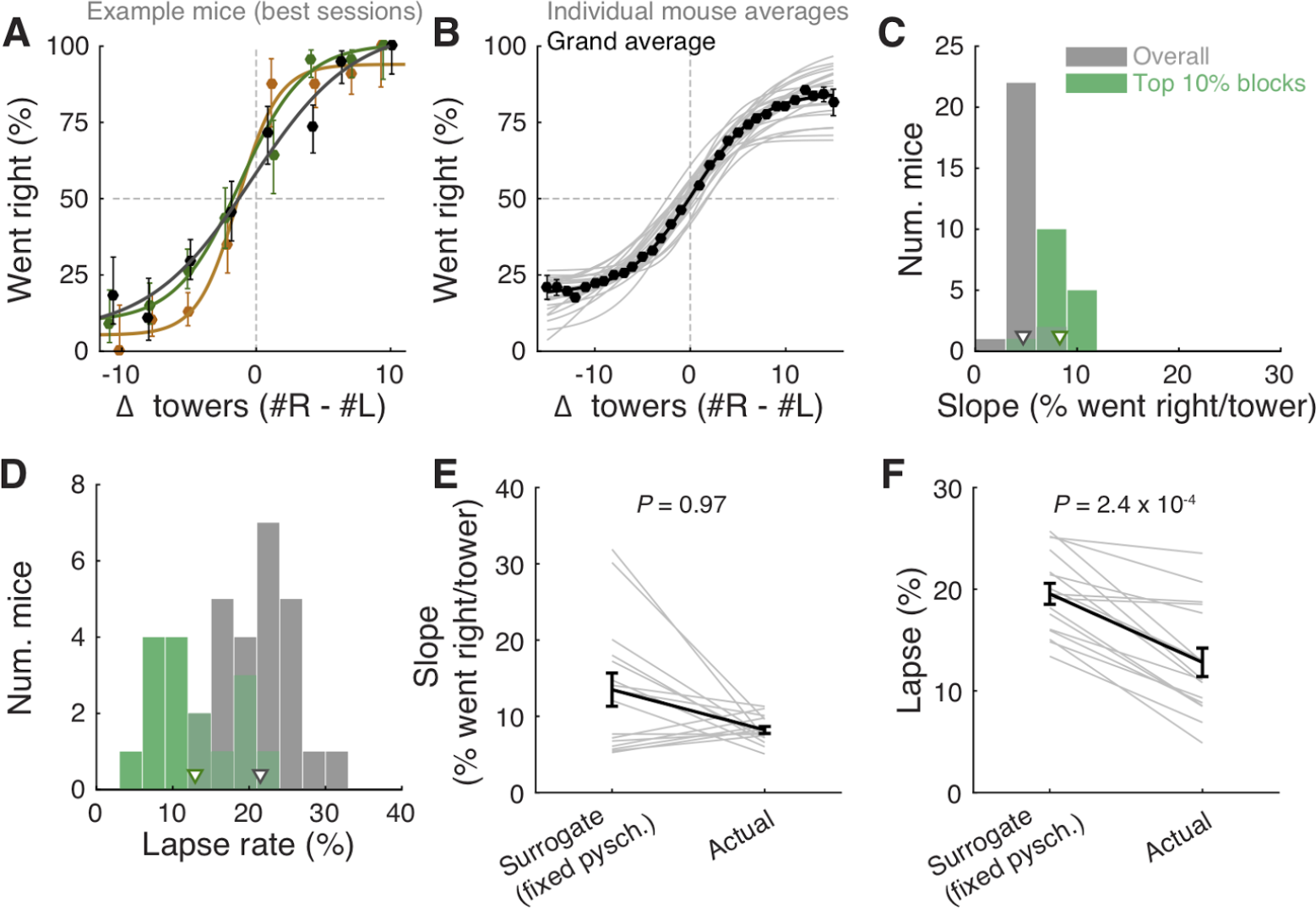
Performance of the accumulating-towers task. **(A)** Best-session example psychometric functions from three mice. Circles: data points, lines: sigmoidal function fits, error bars: binomial confidence intervals. **(B)** Overall psychometric functions across the population.

Thin gray lines: sigmoidal function fits for all mice with at least 1000 trials (n = 25). Black circles and line: psychometric function with sigmoidal fit for aggregate data (metamouse, n = 135,824 trials). Error bars: binomial confidence intervals. **(C)** Distribution of slope of the psychometric function for the individual mice shown in B, pooling all data (gray) or selecting the top 10% of blocks for each animal (with at least 300 remaining trials after this selection, n = 16), as defined by average performance. Arrowheads: mean. **(D)** Distribution of lapse rates for the mice shown in B, defined as the average error rate for trials where |#R – #L| ≥ 10. Conventions as in C. **(E)** Comparison between psychometric slopes obtained for the top 10% of blocks in a surrogate dataset sampled from a fixed psychometric curve vs. the actual data. Thin gray lines: individual mice, black lines: average, error bars: ± SEM. **(F)** Comparison of lapse rates between the surrogate and actual data sets. Conventions as in E.

We also wondered what impact within-session fluctuations in performance had in the measured psychometric functions. We calculated ongoing performance using a sliding gaussian window and recalculated psychometric functions after excluding low-performance bouts (i.e. consecutive trials with performance below several different thresholds). Excluding these trial bouts yielded sharper psychometric curves with lower lapse rates (**Supplementary Figure 3B**). While these criteria were not used for any other analyses in this work, they illustrate how more stringent trial selection criteria may be applied depending on data analysis needs.

Behavioral performance in the accumulating-towers task was thus on average comparable to that seen in rats doing an analogous task (Scott et al., 2015), consistent with the finding that mice and rats perform similarly in several perceptual decision-making tasks (Jaramillo and Zador, 2014; Mayrhofer et al., 2013). Additionally, mice of different strains did not show any statistically significant differences in a variety of performance indicators, except for running speed (**Supplementary Figure 4**). Note, however, that strain comparison was not the main goal of this study, and sample sizes and the specific chosen strains were a function of data available from separate unpublished, ongoing studies. We thus lacked the appropriate sample size to detect small differences in behavior. This caveat notwithstanding, we were able to train mice from all tested strains on the task.

### Mice use multiple evidence pulses from the entire cue region

We next sought to determine whether mice solve the task by using towers from the entire cue period. For each mouse we performed a logistic regression analysis to predict choice using the amount of net evidence in each of 5 equally spaced bins spanning the 200-cm cue region. We observed a variety of shapes in the curves given by the different spatial weights in the model (**Figures 3A, B**): while some mice had fairly flat curves, suggesting spatially homogeneous accumulation of evidence (**Figure 3A**), others had curves with higher coefficients in the beginning of the maze, indicating primacy effects, and a minority had higher coefficients in the later spatial bins, suggesting recency effects (**Figure 3B**). To better quantify this, we computed a weight decay ratio between the average weight in the two last and two first bins, such that numbers smaller than one indicate primacy, and estimated statistical significance of individual animals with a shuffling procedure (**Figure 3C**, see **Materials and Methods**). Across the population, we obtained an average ratio of 0.73 ± 0.06 (mean ± SEM), significantly different than one (*P* = 1.3 × 10^−4^, two-sided t-test). Furthermore, 10/25 mice had indices that were significantly smaller than one. Next, to further quantify the contribution of towers from different portions of the cue period, we calculated the percentage of trials containing cues on the non-rewarded side (minority cues) in the different spatial bins, separately for correct and error trials (**Figures 3D, E**). The overall magnitude of this percentage was significantly different between correct and error trials (*F*_1,24_ = 381.75, *P* = 5.73 × 10^−50^), as expected because trials with a higher density of minority cues are more difficult by design. Unlike what was observed in a different evidence-based navigation task (Morcos and Harvey, 2016), the distribution of trials with minority cues did not vary significantly as a function of position (*F*1,4 = 0.15, *P* = 0.96, 2-way repeated-measures ANOVA). On the whole, these analyses suggest that the mice take into account evidence from the entire cue period, on average slightly overweighting earlier evidence. This is consistent with findings from both humans and monkeys performing pulse-based evidence accumulation tasks (Bronfman et al., 2016; Kiani et al., 2008; Tsetsos et al., 2012).

**Figure 3.**
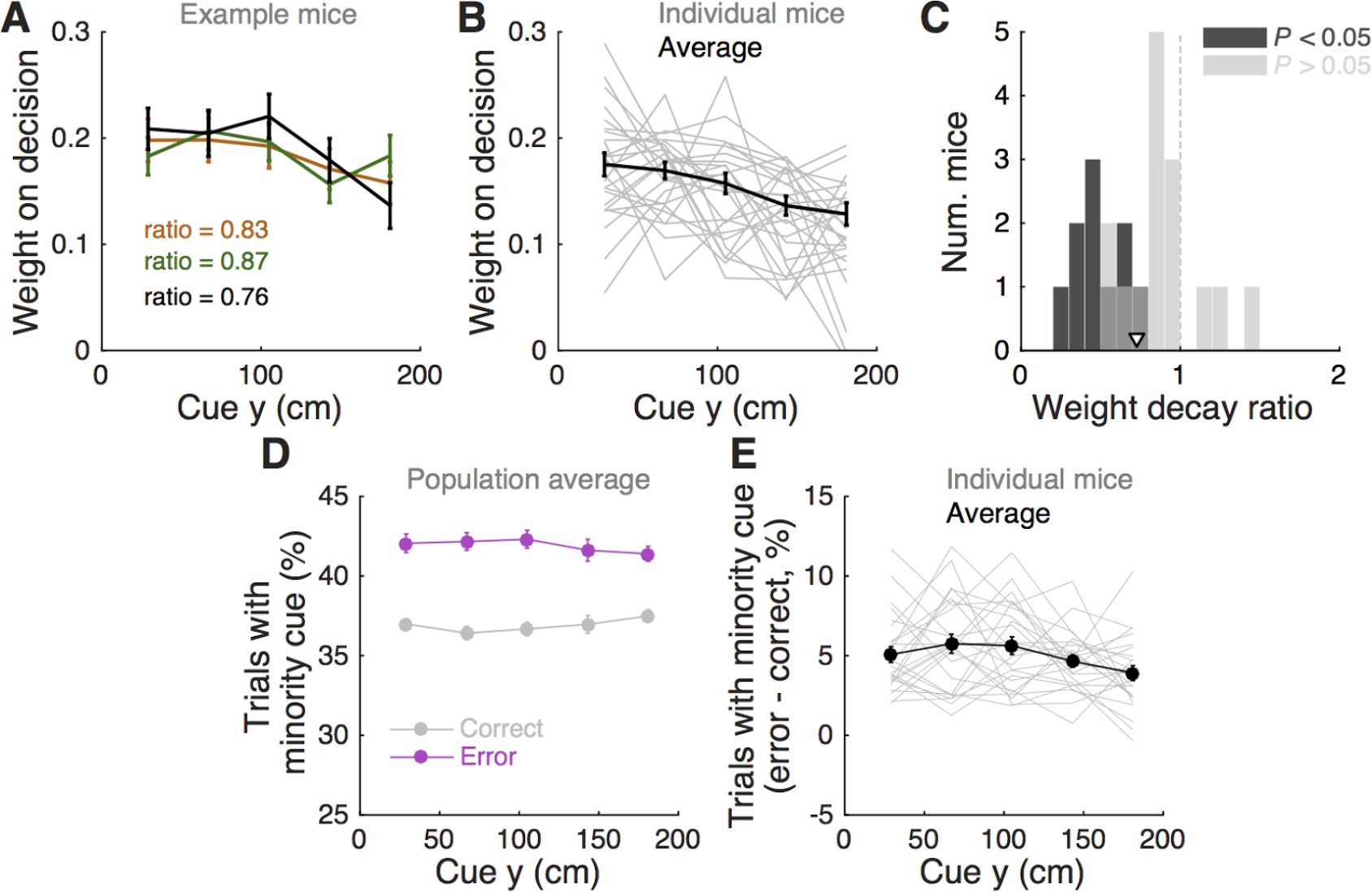
Mice use cues from the entire cue region. **(A)** Example logistic regression for three mice. In this analysis, net evidence (#R – #L) in each of five spatial bins is used to predict the mouse's final decision to turn left or right. Notice fairly flat shapes, suggesting that mice take into account evidence from all parts of the cue period. **(B)** Logistic regression coefficients for all mice with at least 1000 trials (thin gray lines, n = 25), along with average coefficients across the population (thick black line). Error bars, ± SEM. **(C)** Distribution of weight decay ratios for the mice shown in B, defined as the average of coefficients in the last two bins divided by the average of the coefficients in the first two bins. Dark gray: mice with significantly non-flat logistic regression weight curves (*P* < 0.05), light gray: mice with flat curves (*P*; 0.05). Arrowhead: mean. **(D)** Average percentage of trials containing at least one minority cue in each binned cue region position for correct (black) and error trials (magenta). Error bars, ± SEM. **(E)** Difference between the percentage of trials containing at least one minority cue in each binned cue region position between correct and error trials, shown for each individual mouse (thin gray lines), and the average across mice (thick black line). Error bars, ± SEM

In theory, it is possible that some of the aforedescribed findings could be obtained if the mice were selecting (one) random tower(s) in different trials, or, less likely, employing more degenerate strategies that do not rely on sensory evidence at all. To test for these possibilities, we built models that implemented such strategies and compared them against models containing spatially binned evidence terms (**Figure 4**). First, we built a trial-history-only model that only contains previous choice and reward terms, i.e. no evidence is used for the decision (see **Materials and Methods**). This modeled the population of mice more poorly, in terms of having significantly worse cross-validated goodness-of-fit (model information index, *MI*) compared to a model that also has sensory evidence terms (*P* = 8.9 × 10^−5^, n = 20, signed rank test, **Figure 4A**). We next assessed a model in which the mouse uses exactly one randomly selected tower per trial to make a decision. Again, this model had significantly worse *MI* than a model where evidence is used for the decision (*P* = 4.1 × 10^−4^, n = 20, paired t-test, **Figure 4A**; both models also have left/right lapse parameters). Moreover, we reasoned that, in this scenario, for a fixed number of total towers (#R + #L) behavioral performance should vary linearly with #R – #L (Morcos and Harvey, 2016), whereas if the mouse uses multiple cues the psychometric curve should deviate from linearity. The data supported the latter: the experimentally obtained psychometric curve was significantly different than the line predicted by the one-random-tower hypothesis (**Figure 4C**, *P* < 0.01, shuffling test, see **Supplementary Materials and Methods**). Additionally, we reasoned that the one-random-tower hypothesis predicted that, for trials without any minority cues (i.e. no towers on the non-rewarded side), performance should not vary as a function of the number of towers, since any randomly selected tower would lead to a correct decision. This, however, is not what we observed. When we compared trials with fewer than 5 towers to trials with more than 9 towers (all on the rewarded side), performance was significantly higher in the latter case for all mice with sufficient trials for this analysis (*P* < 0.001, signed rank test, not shown). Finally, we implemented other models in which the mice adopt other trivial strategies, namely making a choice based on the first tower, last tower, and 3, 5 or 7 random towers (see **Materials and Methods**). The spatial bin logistic regression model (**Figure 3**) significantly outperformed all five alternative models at the population level (data not shown, n = 20, *P* < 0.01, paired difference tests with false discovery rate correction, see **Materials and Methods**)..

**Figure 4.**
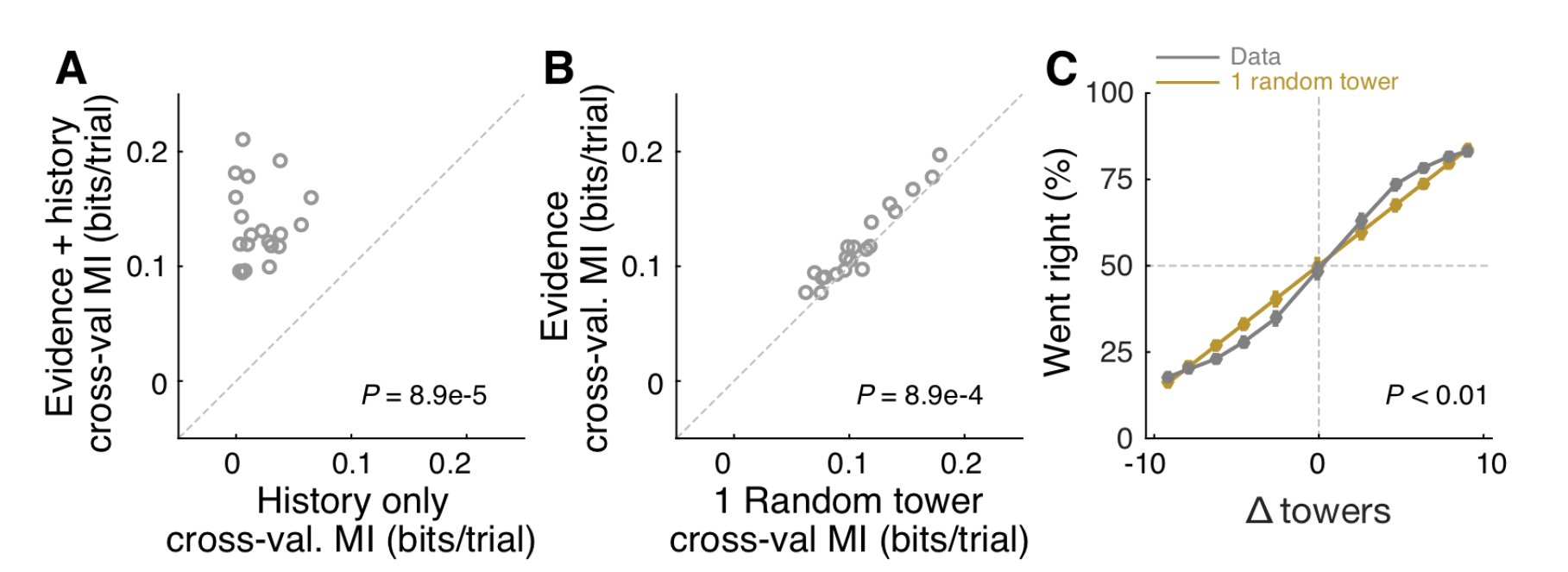
Mice rely on multiple cues to perform the task. **(A)** Comparison of cross-validated prediction performance of a model containing both trial history and spatially binned evidence (**Figure 7H**) and one containing only trial history terms (n = 20 mice). MI: model information. **(B)** Comparison of cross-validated prediction performance of a model in which the mouse makes a decision based on a single random tower and one with spatially binned evidence (no history) (n = 20 mice). **(C)** Psychometric curves for the actual data and a model that chooses from each trial 1 of the presented cues (randomly) and bases the trial choice on the identity of that cue. Data is aggregated across mice for trials where the total number of cues (#R+#L) is equal to 12. In the scenario where #R + #L is fixed, we expect the “1 random cue” model’s performance to be linear with #R – #L (as is borne out in the figure). In contrast, if mice used multiple cues the psychometric curve should be different from a line. The psychometric curve for the actual data (blue) is significantly different from that predicted by the ‘1 random cue’ model (*P* < 0.001, shuffle test, see **Supplementary Materials and Methods**).

### Performance is affected by the number of cues but not trial duration

Having thus established that mice accumulate multiple pulses of evidence from the whole cue period, we next sought to quantify in more detail how the number of towers and cue or delay period duration affected performance. For trials with similar difficulty (same |Δ| towers), we plotted percent correct performance as a function of the effective duration of the cue and delay periods, and noticed no apparent dependence (**Figures 5A, D**). Conversely, when we plotted performance as a function of the total number of towers (#R + #L) for different values of |Δ|, we observed a consistent decrease in performance with increasing numbers of towers (**Figure 5B**), similar to previous findings in the rat (Scott et al., 2015). To quantify how |Δ|, #R + #L and duration each influence performance, we fitted a linear model using these three quantities as predictors (see **Materials and Methods** for details). The largest contribution to performance was given by |Δ| (*P* = 3.1 × 10^−46^, t statistic for regression coefficients), and total number of towers had a significant negative coefficient (*P* = 2.4 × 10^−11^), whereas the cue duration coefficient was not significantly different than zero (*P* = 0.64)(**Figure 5C**).

**Figure 5.**
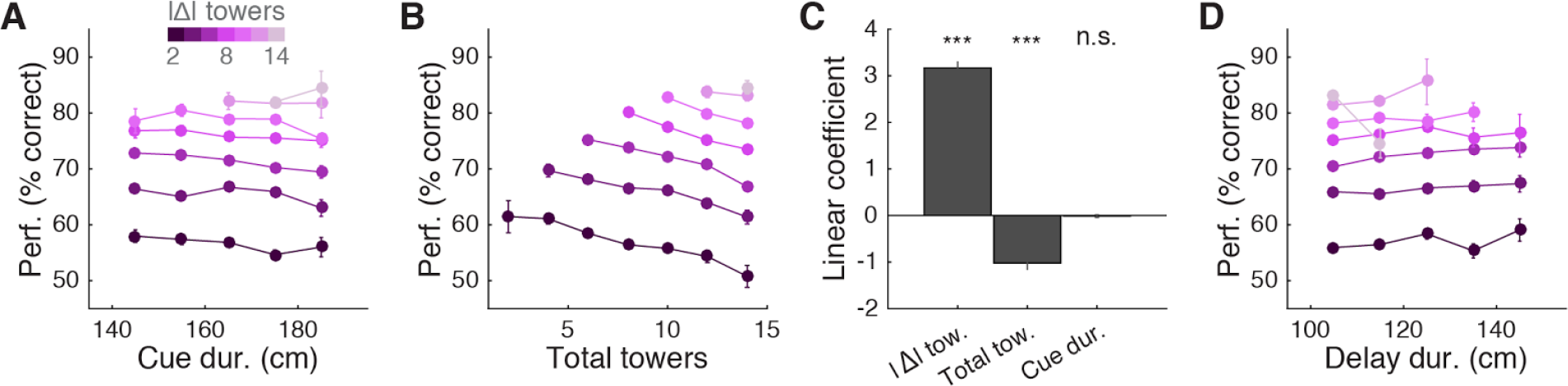
Behavioral performance decreases with increasing total number of towers, but not duration of cue period. **(A)** Overall performance as a function of effective cue period duration in space, for various subsets of trials with different absolute differences between tower counts (|Δ|, color code). Effective duration is defined as the position of the last viewed tower minus the position of the first tower. Error bars: binomial confidence intervals. **(B)** Overall performance as a function of the total number of towers (#R + #L), for subsets of trials with different |Δ|. Conventions as in (A). **(C)** Best-fit coefficients from a linear regression model predicting performance as a weighted combination of |Δ| towers, total towers, and effective cue period duration. The data is the mean-subtracted performance averaged across mice. Error bars: standard error for each parameter. Significance was calculated from parameter *t*-statistics. (**D**) Overall performance as a function of effective delay period duration in space for subsets of trials with different |Δ|. Error bars: binomial confidence intervals. Conventions as in (A).

A potential explanation for the findings described above is the existence of one or more sources of noise that grow proportionally with the number of visual pulses presented in the trial, and that the noise generated by these sources is greater than stimuli-independent noise sources, such as time-dependent accumulation (diffusion) noise in a drift-diffusion model (DDM) framework (Brunton et al., 2013; Scott et al., 2015). Potential sources of stimulus-dependent noise are many, and include noise in stimulus presentation and/or processing, which adds noise independently with each pulse (Brunton et al., 2013; Smith and Ratcliff, 2004), or noise that scales non-linearly with the total amount of pulses (Fechner, 1860; Scott et al., 2015).

To further investigate this, we fitted the data using two different models. First, to estimate the magnitude of different noise sources, we employed a DDM developed by Brunton et al. (Brunton et al., 2013), which models a latent decision variable as a function of memory leak (*λ*), a sticky accumulation bound, and three sources of noise: diffusion, stimulus and initial value of the accumulator (in addition to four other parameters, see **Materials and Methods**, **Supplementary Figure 5**). Consistent with findings in rats (Brunton et al., 2013; Scott et al., 2015), we found that sensory noise was the dominant source of noise for the majority of mice (**Figures 6A,B** and **Supplementary Figure 5**, across the population: *P* = 7.0 × 10^−4^, t test, and 5.9 × 10^−4^, signed rank test, respectively for *σ^2^_s_* vs. *σ^2^_i_* and *σ^2^_s_* vs. *σ^2^_a_*, 8/20 and 9/20 mice had significantly higher *σ^2^_s_* compared to *σ^2^_i_* and *σ^2^_a_*, respectively, based on proportions of cross-validation runs). Also consistent with the previous studies, we found memory leaks close to zero (**Figure 6C**)(*λ* = 0.03 ± 0.09 m-1, mean ± SEM, *P* = 0.78, two-sided t-test vs. zero, only four animals had *λ* values that were statistically different from zero). These results are also consistent with our analyses in **Figure 5**, where neither the duration over which the cues are presented nor the effective delay interval after the last cue are significant factors (beyond #R and #L). Like other DDMs, the Brunton et al. model assumes that each pulse of evidence is associated with independent Gaussian noise, which results in linear scaling of the total variance with increasing number of pulses. This assumption, however, has been shown not to hold for an analogous visual pulse accumulation task in the rat or the acoustic version of Brunton et al. (Scott et al., 2015). Instead, the standard deviation of the perceived evidence (i.e., not the variance but its squared root) increased linearly with increasing number of pulses, favoring a scalar variability framework (Fechner, 1860; Gallistel and Gelman, 2000). We attempted to quantify this in our data by fitting the same Signal Detection Theory (SDT) model as Scott et al. In this model, each unique tower count is associated with a Gaussian distribution of mean number of towers *µ_T_* and standard deviation *σ_T_*, the latter being the free parameter. The probability of choosing a side is given by the difference in the distributions of left and right tower counts (see **Materials and Methods** for details). For the data aggregated across mice (we failed to obtain low-noise parameter estimates from fits to individual mice), best-fit *σ_T_* grew monotonically with the number of towers *T* (**Figure 6D**). We then fitted two competing two-parameter models to directly test whether scalar variability or linear variance predicted the data better. Again in agreement with the visual and auditory rat tasks (Scott et al., 2015), we found that the scalar variability was, on average, a better model than the alternative, although the results were variable at the level of individual mice (for aggregate data: *P* < 0.003, bootstrapping; 10/25 individual mice had significantly better predictions from the scalar variability model, 2/25 had better linear variance, 13/25 were statistically indistinguishable).

**Figure 6.**
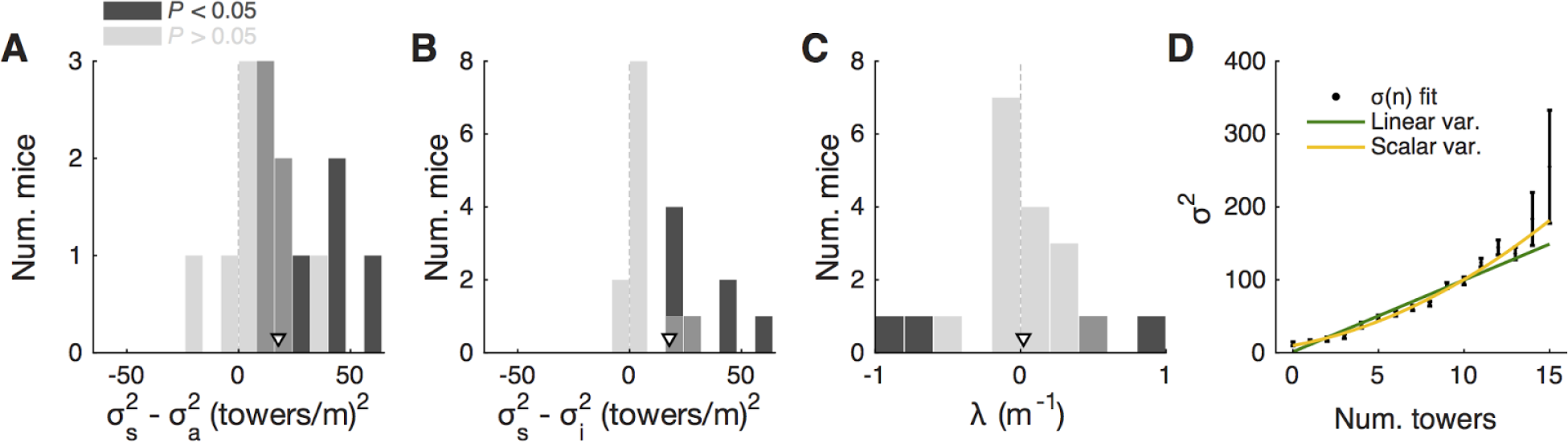
Behavioral performance seems to be limited by cue-dependent noise. **(A)** Distribution across mice (n = 20) of the difference between best-fit sensory and diffusion noise parameters (*σ^2^_s_* and *σ^2^_a_*, respectively) from the Brunton et al. model, color coded according to whether they are significantly different from zero according to 95% confidence intervals determined from cross-validation runs. Arrowhead, population mean. **(B)** Distribution across mice of the difference between best-fit sensory and initial noise parameters (*σ^2^_s_* and *σ^2^_i_*, respectively) from the Brunton et al. model. Conventions as in a. (**C**) Distribution of the memory leak (*λ*) parameter from the Brunton et al. model. Conventions as in a. (**D**) Best-fit parameters for the SDT model (aggregate mouse data). Black data points: parameters from the full model where *σ^2^* is determined separately for each tower count. Yellow line: prediction from the two-parameter scalar variability model. Green line: prediction from the two-parameter linear variance model. Scalar variability yielded significantly better predictions (*P* < 0.003). Error bars: standard deviation from bootstrapping iterations (n = 200).

### Choice is influenced by previous trial history

Having determined that mice are sensitive to the amount of sensory evidence and that they use multiple pulses throughout the maze to make their decision, we next investigated how previous choice and reward history influenced current choice. Rodents have been shown to display behavioral effects of trial history in a variety of task designs (Busse et al., 2011; Narayanan et al., 2013; Pinto and Dan, 2015; Scott et al., 2015). In particular, in two-alternative forced choice tasks in operant conditioning chambers, they are more likely to repeat previously rewarded choices (Busse et al., 2011; Scott et al., 2015). We were therefore surprised to uncover the opposite pattern of trial history, albeit of small magnitude: on average, the mice were more likely to go to the opposite arm to a previously rewarded one, and to repeat an unrewarded choice (**Figures 7A, B**). This behavior is potentially reminiscent of the well-documented tendency to spontaneously alternate arms when mice explore a (physical) T-maze (Lalonde, 2002). To better quantify this effect, we defined the alternation bias for each mouse as the mean-subtracted percentage of trials in which they chose the arm opposite to their previous choice (see **Materials and Methods** for details). Post-error and post-reward biases did not significantly differ in magnitude (**Figure 7C**, *P* = 0.48, two-sided paired *t*-test), although only post-error biases were significantly different than zero (*P* = 0.006 and 0.11 for post-error and post-reward, respectively, two-sided *t*-test). There was no correlation between the magnitude of post-reward and post-error biases across mice (*r* = –0.28, *P* = 0.25, Pearson's correlation, analysis not shown). Post-error biases also had a longer time scale than post-reward, going at least five trials in the past (**Figure 7D**). Note that for the analysis **Figure 7D** a long-lasting negative bias with respect to trial zero indicates higher probability of going to same arm over consecutive trials. In other words, this would indicate the presence of choice perseveration bouts, particularly following an error trial. To directly assess this, we calculated the alternation bias selecting trials with consecutive rewards or errors in the same arm (**Figure 7E**). We noted an increase in the magnitude of negative bias with increasing numbers of consecutive erroneous visits to the same arm, as expected from the interaction between choice perseveration and our debiasing algorithm. For example, a mouse that perseverates in going left with little regard to the evidence will cause the debiasing algorithm to sample more right-rewarded trials, increasing the fraction of consecutive left-choice, right-rewarded trials. To estimate how these perseveration bouts affected overall performance, we recomputed the aggregate psychometric curve after removing trial bouts in which the mice made at least three consecutive identical choices. Applying this additional trial selection criterion resulted in little performance improvement (**Figure 7F**, and changing the criterion to more trials did not qualitatively change the results).

**Figure 7.**
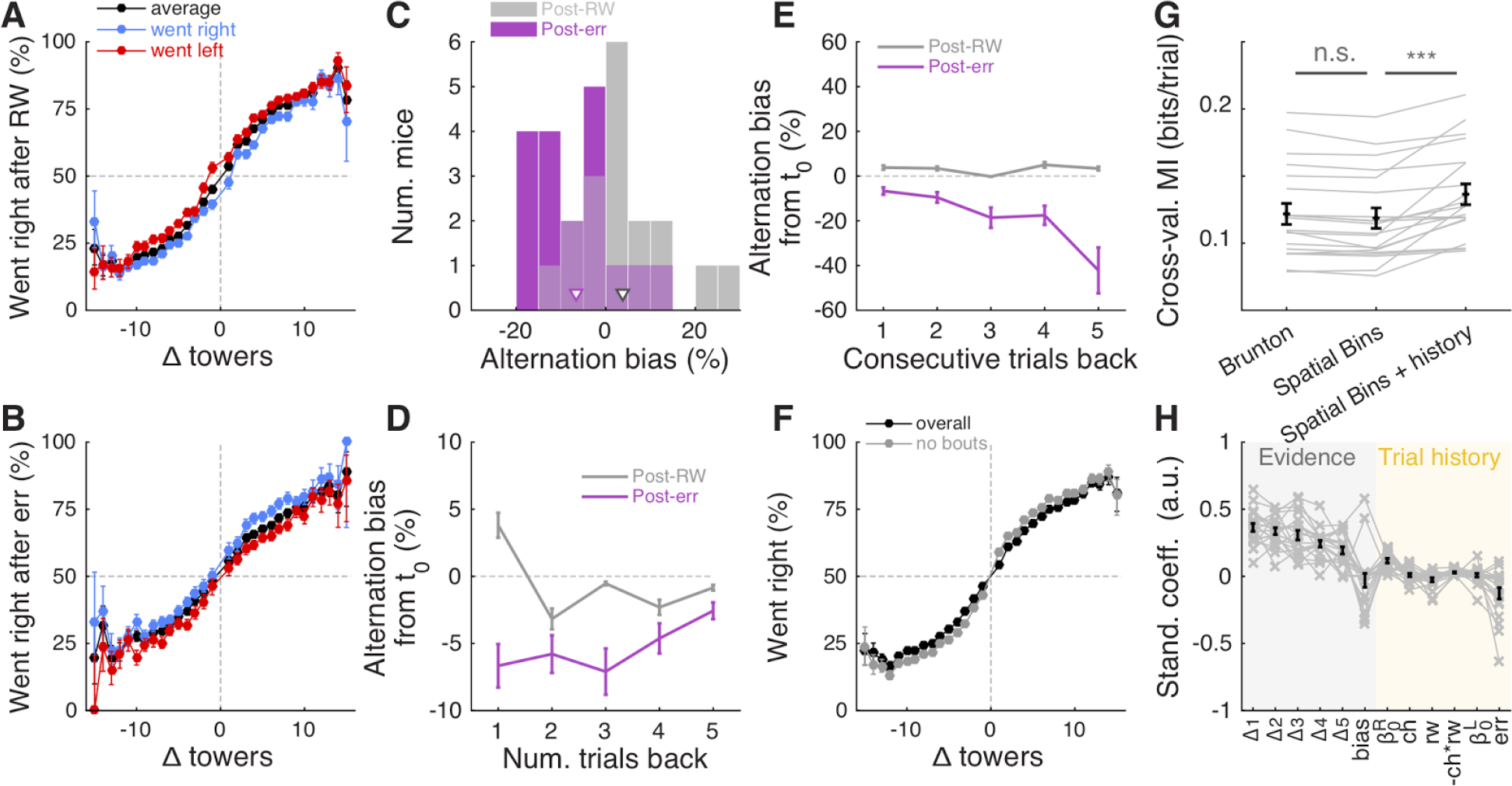
Choice is moderately influenced by previous trial history. **(A)** Psychometric curves for aggregate data (metamouse) divided according to previous choice in rewarded trials. Black: average post-reward curve, blue: psychometric curve for trials following rewarded right choices, red: psychometric curve for trials following rewarded left choices. Error bars: binomial confidence intervals. **(B)** Psychometric curves divided according to previous choice in error trials. Conventions as in (A). **(C)** Distribution of alternation bias after reward (gray) or error (magenta) trials. Arrowheads: population mean. **(D)** Magnitude of alternation bias calculated for 1 – 5 trials after a choice, separately for rewarded and unrewarded trials. Error bars: ± SEM across mice (n = 18 with at least 1,000 trials after removing trials with fewer than 5 history trials). **(E)** Magnitude of alternation bias calculated for 1 – 5 trials after identical rewarded or unrewarded choices. Error bars: ± SEM. **(F)** Psychometric curves for aggregate data (metamouse) with the trial selection adopted throughout the paper (black) and adopting an additional criterion to exclude at least 3 consecutive-choice trials (gray). Error bars: binomial confidence intervals. **(G)** Comparison of cross-validated model prediction performance for the Brunton et al. DDM, the spatial-bin logistic regression, and the latter plus trial history terms. Thin gray lines: individual mice, black lines and error bars: mean ± SEM. *** *P* < 0.001. MI: model information index. **(H)** Best-fit standardized coefficients for the spatial bins model with trial history terms. Thin gray lines: individual mice, thick black lines: population mean, error bars: ± SEM.

Thus, choice and reward history impacted present choice in the accumulating-towers task. To provide a more complete description of the behavior, we added trial history to our behavioral models. First, however, we compared the performance of the Brunton et al. DDM to the logistic regression model in which choice is a weighted function of the net amount of sensory evidence in different spatial bins (**Figures 2A, B**). This latter heuristic model performed as well as the Brunton DDM in cross-validated datasets across the population (**Figure 7G**, *P* = 0.58, n = 20, Tukey's post-hoc test after a one-way ANOVA with repeated measures for the three models in the figure; *P* = 2.4 × 10^−6^ for main effect of model type). We therefore added trial history to the logistic regression model (**Figure 7H**), adding terms to account for both the observed vertical shifts in the psychometric curves and the decrease in psychometric slope following errors (**Figures 7A, B**, see **Materials and Methods** for details). As expected, doing so significantly increased model performance (**Figure 7G**, *P* = 9.9 × 10^−5^, Tukey's post-hoc test). Note that this model has the underlying assumption that the mice adopt a spatial strategy, i.e. that weights are assigned to net evidence in different segments of the nominal cue region, regardless of how many towers the mice have seen before reaching that segment. An alternative hypothesis is that mice weight towers according to the order in which they occur. In this scenario, the first tower in a trial would have the same impact on the mouse's decision, whether it occurred on the first or nth spatial segment. To test for this possibility, we constructed another logistic regression model in which towers are ranked (in bins of three) according to their ascending order of occurrence (**Supplementary Figure 6**). The model, which confirmed the predominance of primacy effects on the behavior (i.e. earlier cues have more weight on the decision), performed marginally better than its spatial counterpart, with a trend towards significantly better cross-validated predictions (*P* = 0.07, n = 20, two-sided paired *t*-test, both cases had the same trial history parameters). This suggests that an internal evidence weighting function that depends on a running numerosity explains behavior at least as well as one that weighs evidence based on a tower’s spatial position.

### Mice display fairly stereotyped running patterns

Lastly, we turned to the analysis of the mice's running patterns as they navigated the maze. We characterized the time course of movements that the mice made, as well as how these are modulated by choice and evidence, as this navigational behavior is presumably reflective of the ongoing decision process in a given trial. Secondly, we quantified the motor skill element of our task, as it generally adds to the difficulty and may specifically contribute to the observed lapse rates of the mice.

Inspection of single-trial speed vs. maze position (trial time) traces suggested that mice run at fairly stereotyped speeds in different portions of the maze (**Supplementary Figure 7A**). Average running speed in the maze stem across the population was 61.1 ± 2.4 cm/s (**Supplementary Figure 7B**, mean ± SEM, range: 44.2 – 92.9), translating into a nominal cue period duration of 3.4 ± 0.1 s (mean ± SEM, range: 2.2 – 4.5). Given the broad distribution of speeds we observed across mice (**Supplementary Figures 7C, D)**, we next wondered whether there was any systematic relationship between running speed and performance across the population. Indeed, we found a significant correlation between these two indicators, averaged across all sessions (**Supplementary Figure 7E**, *r* = 0.48, *P* = 0.02, Pearson's correlation). In other words, faster mice tended to perform better. This relationship, however, did not in general hold within individual mice in a session-by-session analysis (**Supplementary Figure 7F**, *r* = 0.06 ± 0.05, mean ± SEM).

We also sought to analyze how frequently mice made putative motor errors (**Supplementary Figure 7G**). We found an overall low occurrence of trials with unusual motor events, at an average of 5.6 ± 1.0 % (mean ± SEM, excluding low-speed trials, which are 10% by definition, see **Materials and Methods**). The distribution of these events differed significantly among event types and between correct and error trials (**Supplementary Figure 7G**, 2-way repeated measures ANOVA, *P_event type_* = 2.8 × 10^−38^ *P_trial outcome_* = 2.6 x 10^−5^), with events being more common in error trials. The frequency of unusual motor events did not depend on trial difficulty (2-way repeated measures ANOVA, *P* = 0.25, data not shown). We thus conclude that while motor aspects did contribute to error trials, they cannot fully explain the lapse rates.

We then turned to the analysis of view angle trajectories. Specifically, we first looked for statistical correlations between view angle distributions and the animal’s behavioral strategy, and find that these distributions do on average reflect the eventual choice of the animal, as follows. For each mouse, we calculated the distribution over trials of view angles as a function of y position in the maze, separately for left and right choice trials (**Figure 8A**). We observed diverging distributions with increasing y positions, indicating that the animals progressively turned to their choice side as they ran down the stem of the maze. To better quantify this phenomenon, we built choice decoders that predicted the future choice based on the current view angle at a particular y position along the maze stem (see **Materials and Methods**). At y = 100 cm (half-way through the cue period), average decoding accuracy was 73.1 ± 1.3% (mean ± SEM), whereas at the end of the cue period (y = 200 cm) we could predict choice with an accuracy of 87.3 ± 1.1% (**Figure 8B**). We reasoned that this divergence of view angles during the cue period could be related to the observed primacy effects (i.e. mouse weighting earlier evidence more, **Figure 2** and **Supplementary Figure 6**), prompting us to look for such relationship at the subject level. We thus calculated the correlation between the weight decay ratio (**Figure 2C**) and choice decoding accuracy at y = 100. We found a significantly negative correlation between the two (**Figure 8C**, *r* = -0.71, *P* = 7.6 × 10^−5^, Pearson's correlation), indicating that in fact animals integrating more evenly across the maze also tended to run straighter during the cue period. This finding possibly indicates that they commit later to a particular decision. In fact, the view angle trajectories of mice were on average modulated by the strength of evidence within the trial, with view angles diverging earlier towards the target side (as defined by the eventual choice) for trials with larger magnitudes of |Δ| towers (**Figure 8E**). Although highly variable on a per-trial basis (**Supplementary Figure 7D**), this aspect of the navigational behavior seemed to be sensitive to parameters of the cognitive strategy employed by the mice (weighting of cues vs. space, trial difficulty), and may prove useful for future studies.

Given the above correlations, we considered the possibility that mice may actually circumvent the memory demands of the task by using the view angle throughout the maze as a mnemonic for the side with more evidence. To address this, we looked for timepoint-by-timepoint changes that could reflect such a mnemonic strategy, e.g. always turning by +1° every time a right cue appears. We computed tower-triggered view angles separately for left and right choices, subtracting the average trajectories for each mouse (**Figure 8D**). As a population, mice exhibit no pulsatile nor step-like changes in the view angle for at least 50 cm (~1s) post either left or right cues, although the aforementioned statistical correlation between view angle and choice is visible at 80cm post-cues (for cues at the end of the cue period, this corresponds to the location of the T-maze arms). However, a mnemonic strategy must be carried out on a per-trial basis, and should be evident in the cue-triggered view angle distribution across *trials* (as opposed to across mice). The 1-standard-deviation spread of view angle across trials is more than a factor of 5 times larger than the difference in right/left choice mean view angles for every mouse (**Supplementary Figure 8**), which argues strongly against a stereotyped, memoryless mnemonic of using an “accumulated” view angle to make a choice.

**Figure 8.**
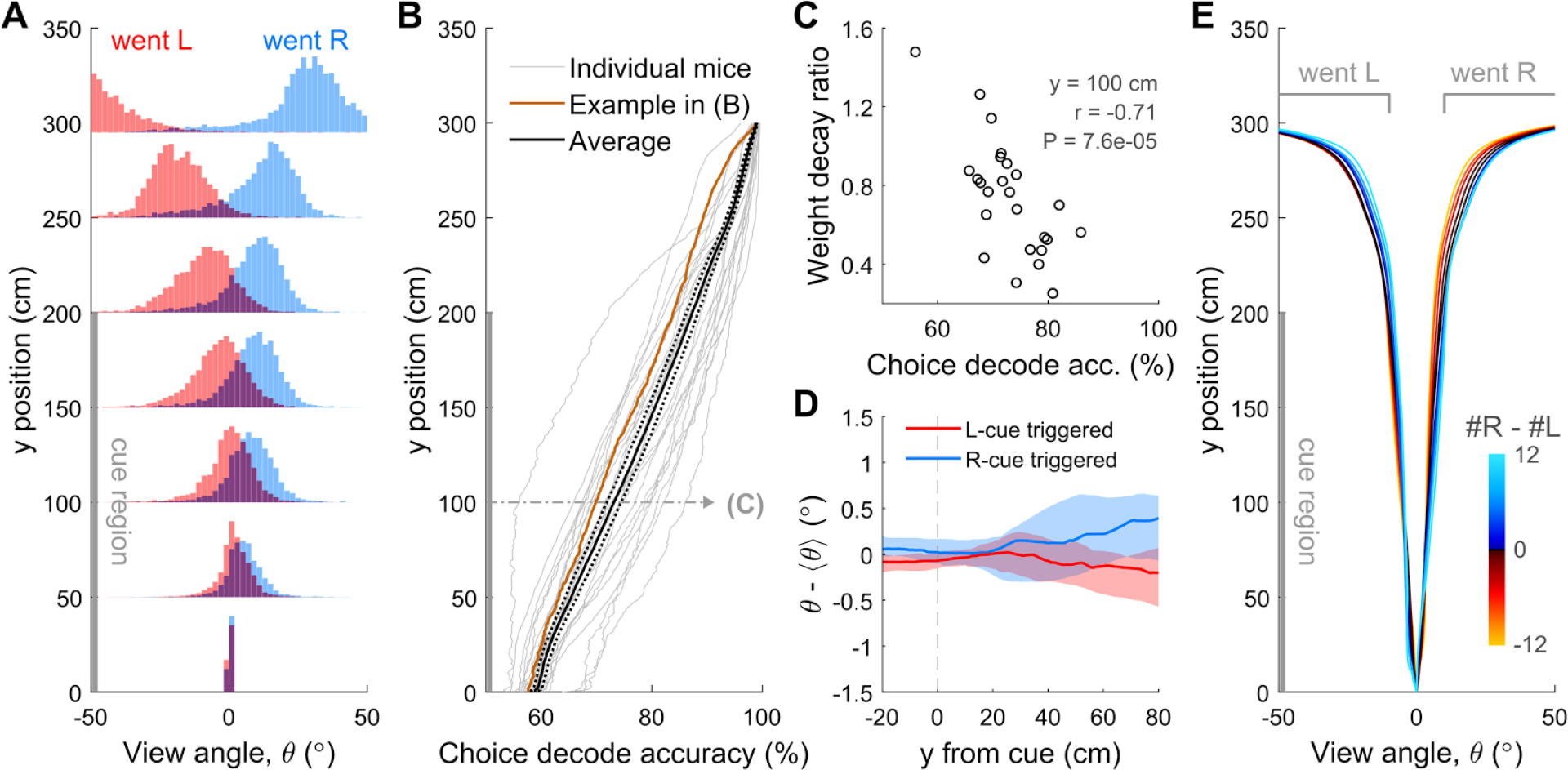
View angle trajectories. **(A)** Distribution of view angles in left and right choice trials (arbitrary units, normalized to equal area for both choice categories) for an example mouse, sampled at several y positions (0, 50, …, 250, 295 cm) along the stem of the T-maze. **(B)** Accuracy of decoding the eventual choice of a given mouse using a threshold on the view angle, evaluated at various y positions along the T-maze. **(C)** Scatter plot across mice of the evidence weight decay ratio (see **Figure 3C**) vs. the choice decoding accuracy evaluated at halfway into the cue region as indicated in **(B)**. **(D)** Cue-triggered change in the view angle θ relative to the average trajectory 〈θ〉 for trials of the same choice. The bands indicate the 1 standard deviation spread across mice, with the thick lines being the median across mice. **(F)** Average view angle for subsets of left/right choice trials with various values of #R – #L (color code). For a given choice, the mean view angle trajectory of individual mice are aligned to the aggregate data (metamouse) before averaging.

## DISCUSSION

We have developed a new virtual navigation-based, pulsed evidence-accumulation task for head-fixed mice, along with tools to quantify their performance and behavioral strategy. We show that mice can gradually accumulate visual evidence in virtual reality over seconds. First, large numbers of mice from several strains can be reproducibly trained in the task (**Figure 2, Supplementary Figures 1–4**). Second, using a combination of analytical and modeling approaches, we also show that mice solve this task by using multiple pulses of evidence across the cue region, although they tend to slightly overweight earlier evidence (**Figures 3, 7** and **Supplementary Figure 6**). Moreover, our analyses suggest that sensory evidence-dependent noise, but not accumulation memory, is an important performance-limiting factor, much like analogous tasks in rats and humans (**Figures 5** and **6**) (Brunton et al., 2013; Scott et al., 2015). An intriguing difference from previous reports was our observation that the mice tended to alternate instead of repeating a previously rewarded choice (**Figure 7**), unlike what has been observed in both rats and mice (Busse et al., 2011; Scott et al., 2015). We speculate that this is related to the mice's tendency to spontaneously alternate their choices in T-mazes (Lalonde, 2002), but note that choice alternation has also been reported in humans performing perceptual decision making tasks, and has been found to be modulated by the magnitude of uncertainty in the previous trial (Urai et al., 2017). Finally, analysis of the mice's virtual navigation trajectories suggested that ongoing behavioral readouts may provide useful proxies for latent cognitive variables (**Figure 8**), although further studies will be needed to explore that possibility in more detail. For example, the slight modulation in view angle trajectories by the amount of sensory evidence could statistically reflect decision confidence (Kepecs et al. 2008), and/or on average different decision times for trials of different difficulty. Additionally, this feature of the behavior could potentially be explored to study changes of mind at the level of single trials (Kiani et al. 2014), by explicitly designing the evidence pulse streams to change their underlying rates at various points in the cue region.

A central aspect of the accumulating-towers task is that decision-making must occur while the animal navigates a (virtual) environment. Despite introducing complexity in the behavioral training and data analyses, we argue that this is a desirable feature. Natural behavior seldom occurs in isolated modules, and is instead dynamic and high-dimensional, and it is precisely these behavioral constraints that are thought to have shaped the evolution of neural circuits (Darwin, 1998; Gomez-Marin et al., 2014; Krakauer et al., 2017). The study of highly reduced decision-making behaviors have allowed the field to make large strides in understanding their underlying neural mechanisms (Brody and Hanks, 2016; Carandini and Churchland, 2013; Gold and Shadlen, 2007). The study of these processes under more complex contexts should yield novel insights into how they are flexibly composed to produce real-world solutions.

A recent study has described a similar visual evidence-accumulation task for mice navigating in VR (Morcos and Harvey, 2016). The accumulating-towers task differs from theirs in a few crucial ways, in terms of stimulus design, task difficulty and apparent strategies adopted by the mice. We used Poisson-distributed, brief pulses of spatially discrete evidence (200 ms, 12-cm separation), which resulted in up to 16 cues on one side (median: 4) and up to 25 cues total (median: 10). Conversely, Morcos and Harvey always had six cues of an optical flow (wallpaper) nature that were four times as long (~800 ms) and occurred in stereotyped positions throughout the stem of the maze. The latter design sampled the same stimulus configurations at high frequencies, which is beneficial for increasing statistical power via averaging. However, we argue that there are complementary advantages to sampling a much larger region of stimulus space with spatially random cues. For example, decorrelating cue locations from space/time allowed us to tease apart the effects of stimulus strength vs. an important aspect of working memory, i.e. retention time (**Figure 5**), while maintaining a quasi-fixed trial duration. Moreover, using brief pulses of sensory evidence gives one the ability to study cue-triggered neural responses (Koay et al., 2016; Scott et al., 2017). This highly heterogeneous design did likely increase task difficulty, which may explain the slightly lower performance we observed compared to the Morcos and Harvey task. Note, however, that we used deliberately liberal trial selection criteria, and that when more stringent criteria were applied we could obtain very high performance sessions (**Figure 2, Supplementary Figure 3**), which might be desirable for neural recording and perturbation experiments. Interestingly, these task design differences led to apparent differences in the strategies that the mice employed. Specifically, the mice in the Morcos and Harvey study displayed more pronounced primacy effects than ours (**Supplementary Figure 9**).

The primacy effects we observed in many of our mice (**Figures 3, 7** and **Supplementary Figure 6**) agree with several other evidence pulse-based tasks in mice, monkeys and humans (Bronfman et al., 2016; Kiani et al., 2008; Ludwig et al., 2005; Odoemene et al., 2017; Tsetsos et al., 2012), but are at odds with findings of temporally even evidence integration in rats performing a high-rate auditory clicks task (Brunton et al., 2013). The reasons behind these differences are a matter of ongoing debate. In particular, it has been argued that primacy in pulsed-based tasks is due to either reaching an accumulator bound (Kiani et al., 2008) or to competition between leaky integrators that mutually inhibit each other (Tsetsos et al., 2012). Interestingly, it has been recently shown that in humans the degree of primacy and even the monotonicity of the evidence weighting curve can change with stimulus duration, which prompted the authors to postulate a dynamic evidence accumulation mechanism (Bronfman et al., 2016). Thus, it is conceivable that different decision-making and integration mechanisms might be at play depending on stimulus and task features (Piet et al., 2017; Uchida et al., 2006). Task design differences could also explain why we did not observe an improvement in performance with increased stimulus durations (**Figure 5**), as might be expected if a diffusion-to-bound-type mechanism is at play. Specifically, our stimulus period durations were longer than when the benefits of prolonged stimulus saturate (Brunton et al., 2013; Gold and Shadlen, 2007; Kiani et al., 2008). On the other hand, the finding that behavior in our task was influenced by the number but not duration of cues is consistent with multiple previous studies of counting in rodents (Mechner 1958; Fernandes and Church 1982; Gallistel and Gelman 2000; Çavdaroğlu and Balc; 2016). Counting is thought to be carried out as a magnitude-estimation process that displays the property of scalar variability, i.e. the noise (standard deviation) in estimates scales linearly with count/magnitude (Fechner 1860; Gallistel and Gelman 2000). Accordingly, following a recent report in rats (Scott et al. 2015), we show through modeling that noise in the mice's estimates of the number of towers in our task scales in a way that is compatible with the phenomenon of scalar variability. We extend previous findings by showing that, in addition to self-generated lever presses (Çavdaroğlu and Balc; 2016), mice can accumulate visual stimuli in the context of a perceptual decision-making task.

In summary, the accumulating-towers task is a valuable behavioral tool to study evidence accumulation and decision-making in mice. The task is conducive to further automation and scaling, and interesting modifications such as designed stimulus sets can be easily incorporated. Most importantly, it is readily integratable with any number of optical or electrophysiological techniques requiring head fixation (Koay et al., 2016; Pinto et al., 2017), allowing us to leverage the comprehensive mouse toolkit in understanding neural mechanisms underlying this important cognitive behavior.

## Author contributions

L.P., S.A.K. and B.E. collected the data; L.P., S.A.K., B.E. and A.M.Y. analyzed the data; S.Y.T., L.P. and B.E. and built the VR systems; S.A.K., B.D., B.E., L.P. developed the task and shaping procedures with guidance from C.D.B., D.W.T. and I.B.W.; L.P., S.A.K., D.W.T. and C.D.B. conceived and wrote the manuscript.

## Acknowledgments

We thank B.B. Scott, J.W. Pillow and A.T. Piet for helpful discussions, and S. Stein for technical assistance.

## Supplementary Materials

### Supplementary Figures, Movie and Table

**Supplementary Figure 1.**
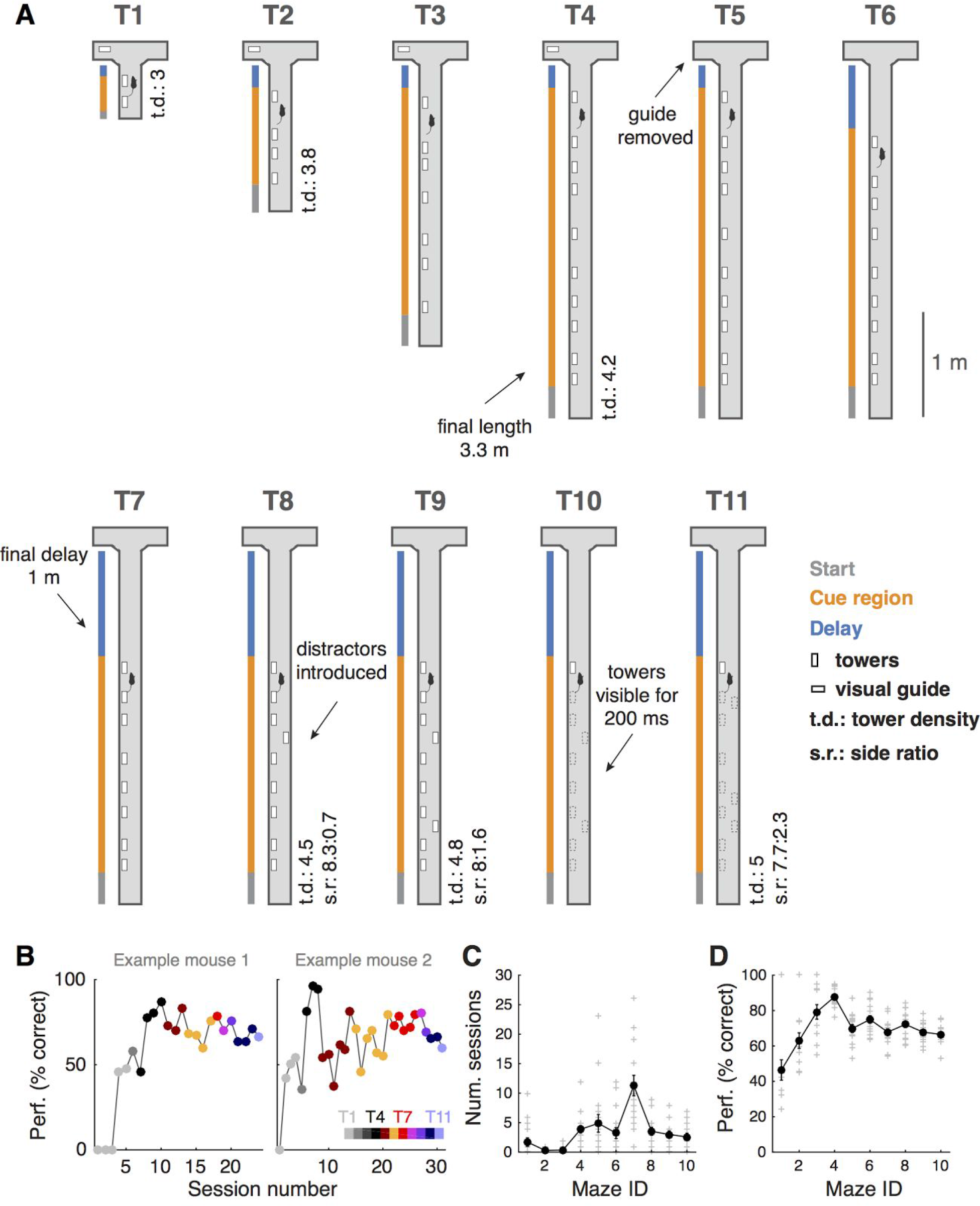
Shaping. **(A)** Schematic illustration of the 10 different shaping mazes (T1 – T10) and the final accumulation maze (T11). **(B)** Progression through shaping stages of two example mice, where each color indicates a different maze according to the colorbar on the bottom right. **(C)** Number of sessions training sessions spent on each shaping stage. Gray crosses: individual mice, black circles: population mean (n = 17), error bars: ± SEM. **(D)** Average overall performance for each shaping stage. Conventions as in (C).

**Supplementary Figure 2.**
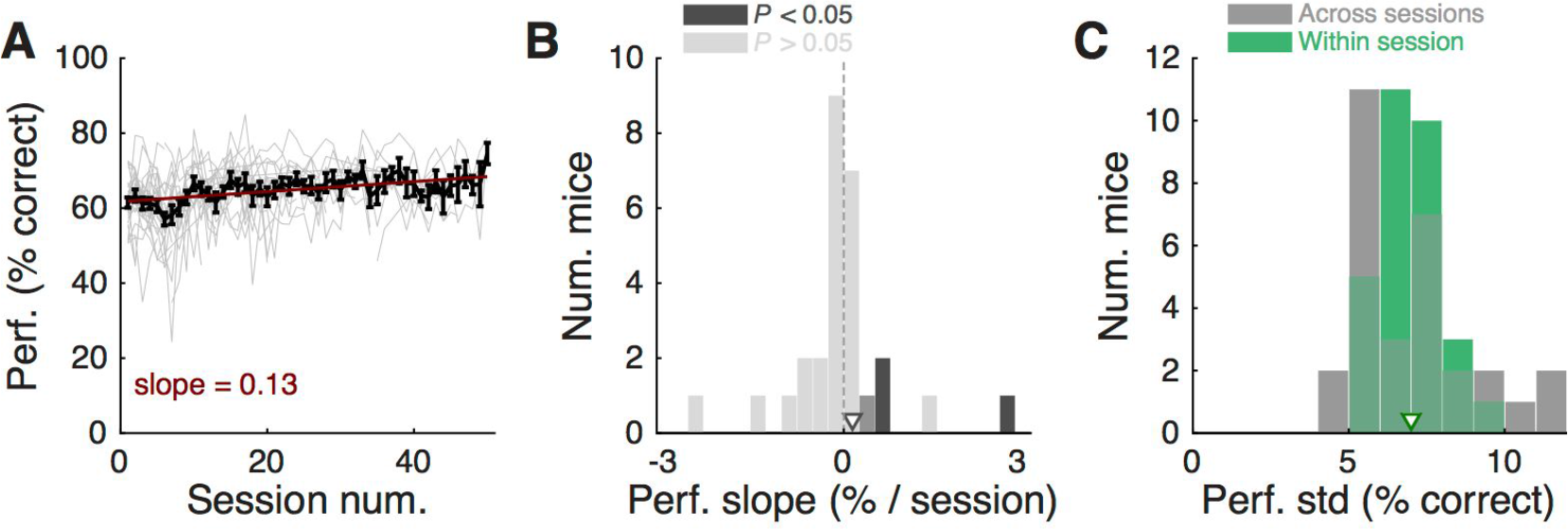
Mice display stable performance over many sessions. **(A)** Overall performance in the final accumulation maze as a function of session number, for mice with at least 5 sessions (n = 30), and selecting all trials regardless of overall performance (i.e. not applying any performance thresholds). Thin gray lines: individual mice, black line: average across mice, error bars: ± SEM. Red line is best linear fit to average data. **(B)** Distribution of slopes extracted from best-fitting lines to performance of each mouse as a function of session number (i.e. thin gray lines in panel A). Bars are color-coded according to whether the slope is significant (dark gray, i.e. its 95% confidence interval does not overlap zero) or not (light gray). Arrowhead indicates population mean. The distribution was not significantly different from zero (*P* = 0.99, signed rank test), indicating stable performance across sessions. **(C)** Distribution of standard deviation of average performance across (gray) and within (green) sessions for the mice. Arrowheads: population means. Within-session standard deviation was calculated using performance over a 40-trial running window.

**Supplementary Figure 3.**
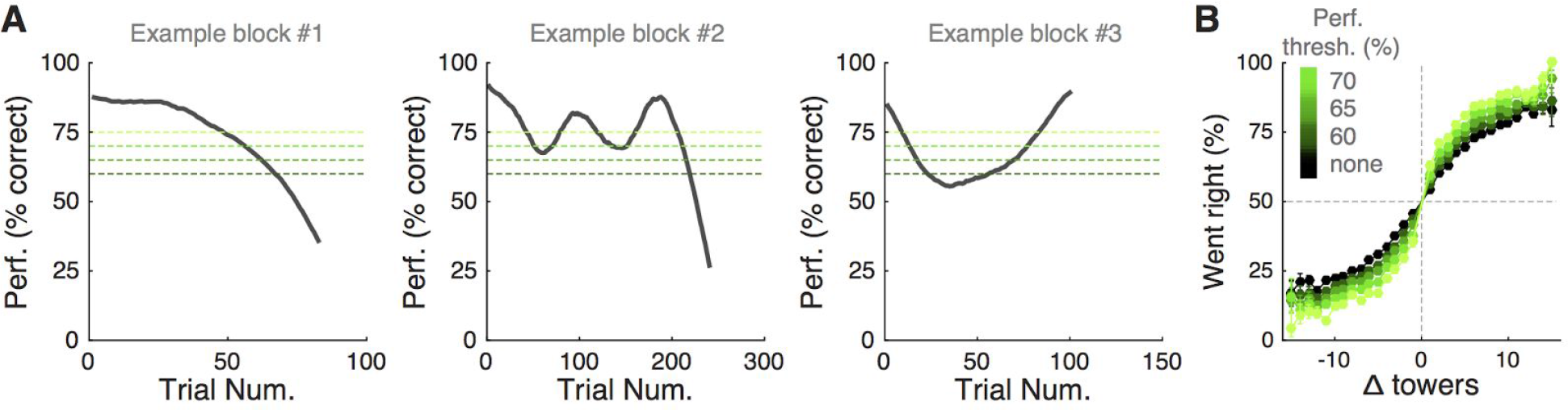
Mice can undergo bouts of high/low performance. **(A)** Three individual examples of consecutive trial blocks on the final accumulation maze, showing performance calculated with a sliding half-Gaussian window (*σ* = 15 trials), plotted as a function of trial number (trial 1 is the first within the block, not necessarily the first in the session). Dotted lines of different shades of green indicate performance thresholds applied in the analysis shown in D. **(B)** Psychometric curves for aggregate data (metamouse), obtain after excluding trials with performance below different thresholds (illustrated in A).

**Supplementary Figure 4.**
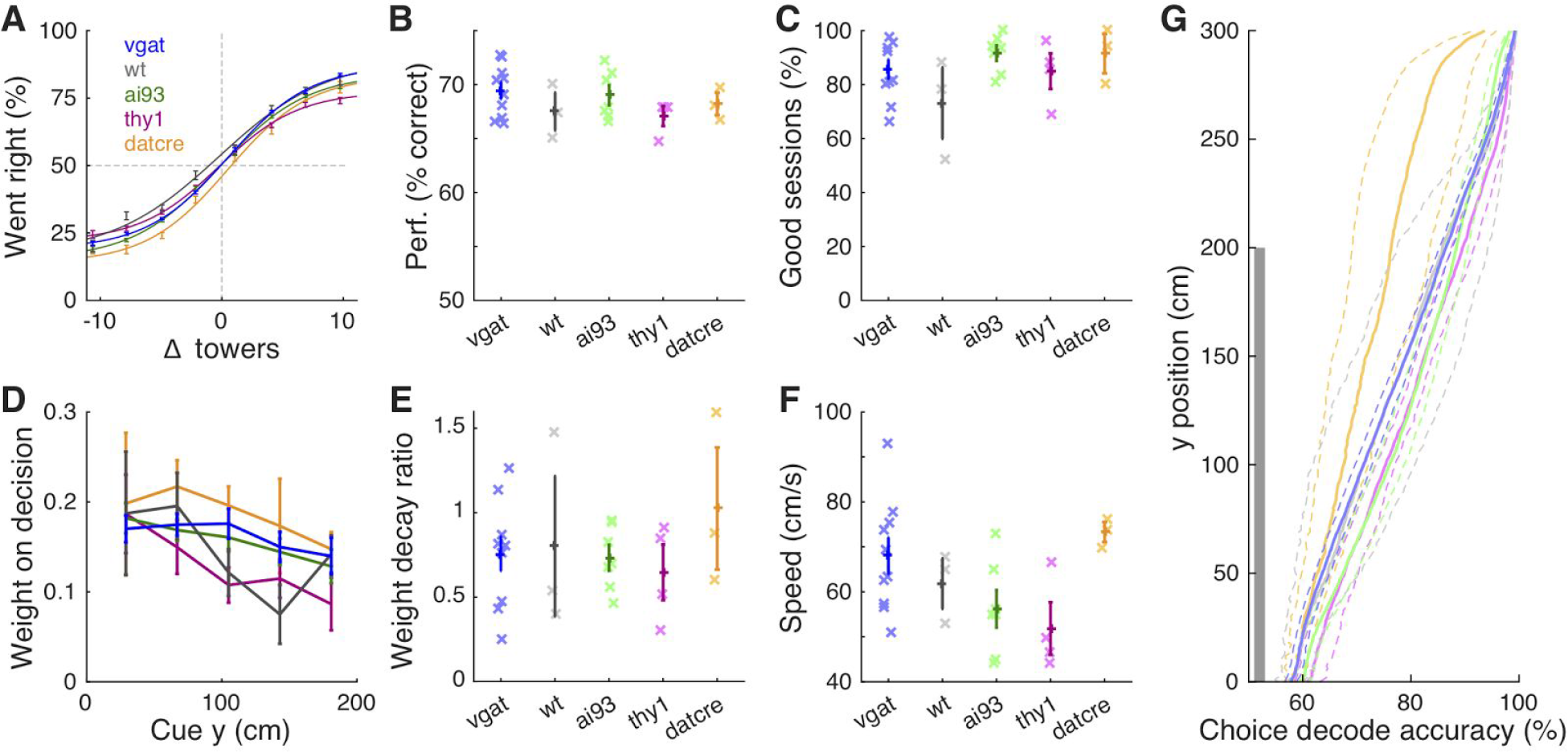
Different mouse strains have comparable performance. **(A)** Psychometric curves for aggregate data (metamouse) divided into five different strains according to the color code on the top left. Error bars: binomial confidence intervals, lines: best sigmoidal function fits. **(B)** Overall performance averaged across blocks with performance over 60% (see Materials and Methods). Crosses: individual animals, error bars: SEM for each mouse strain. **(C)** Percentage of sessions with at least one trial block over our performance threshold (60%). Conventions as in B. **(D)** Logistic regression of choice on net evidence for each spatial bin. **(E)** Weight decay index according to genotype. Conventions as in B. **(F)** Average running speed, conventions as in B. **(G)** Accuracy of decoding choice from view angle as a function of maze position for different strains. For all but one measure above, there was no significant difference between the different strains, (*P* > 0.05, one-way ANOVA)(Decoding accuracy was measured in the cue period, y < 200 cm). The exception was running speed, significantly different between genotypes (P = 0.04).

**Supplementary Figure 5.**
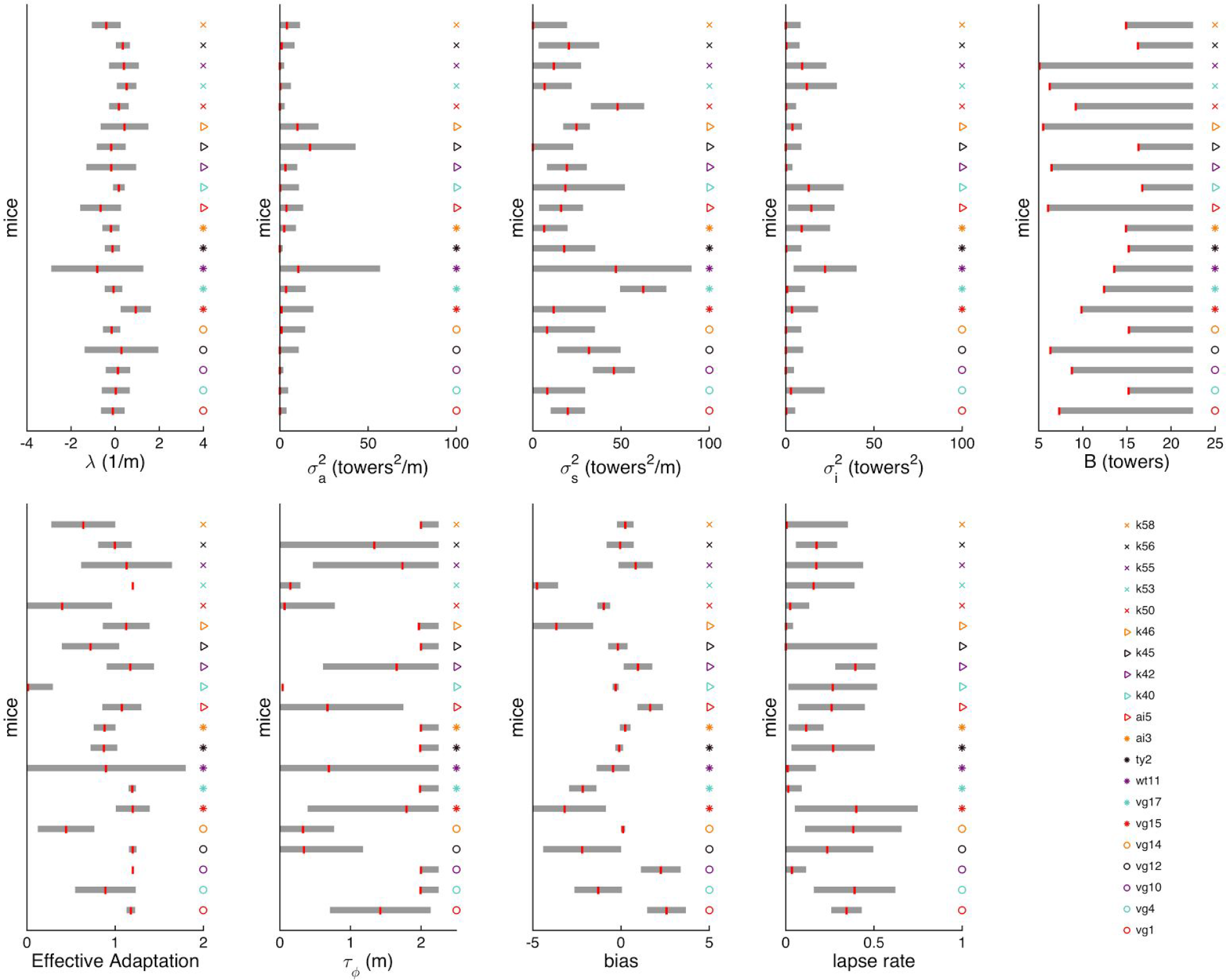
Best-fit parameters for the Brunton et al. model for each mouse. **(A) – (I).** Vertical red bars indicate the median of best-fit parameters across cross-validation runs, gray shadings indicate one standard deviation of the distribution obtained from cross-validation runs. All panels are sorted according to the same mouse order.

**Supplementary Figure 6.**
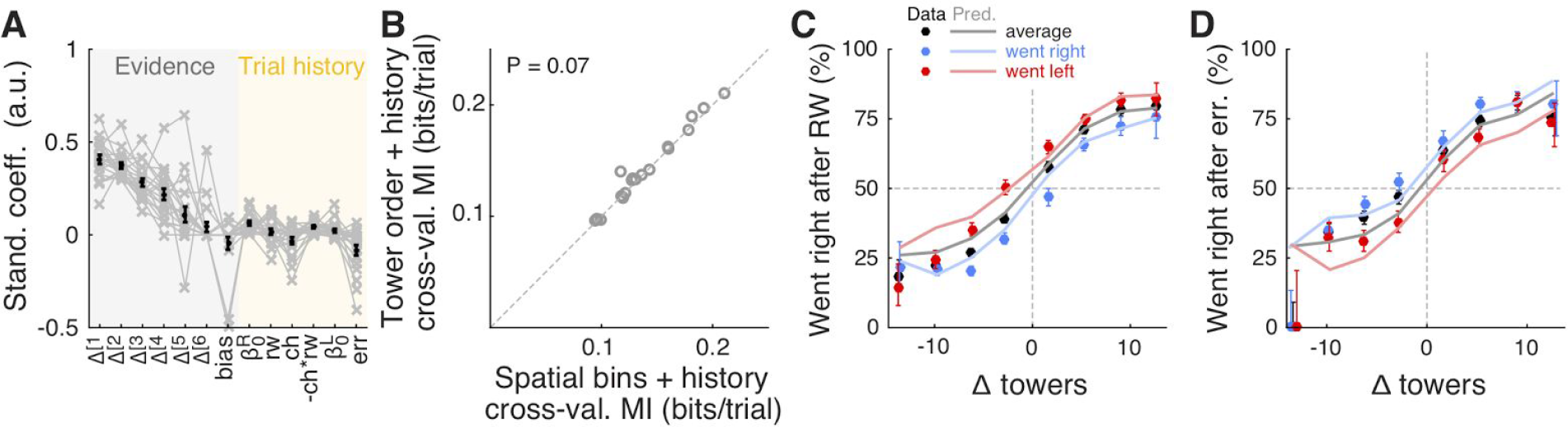
Tower order explains behavior at least as well as position does. **(A)** Best-fit model coefficients for the tower order model with trial history terms. Thin gray lines: individual mice, thick black lines: population mean, error bars: ± SEM. **(B)** Comparison of cross-validated prediction performance of the spatial bins and tower order models, both with trial history (n = 20 mice). MI: model information index. **(C)** Psychometric curve predictions for an example mouse with large trial history effects, divided according to previous choice in rewarded trials. Circles: data, lines: model prediction. Black: average post-reward curve, blue: trials following rewarded right choices, red: trials following rewarded left choices. Error bars: binomial confidence intervals. **(D)** Psychometric curve predictions for the same mouse in (C), divided according to previous choice in error trials. Conventions as in (C).

**Supplementary Figure 7.**
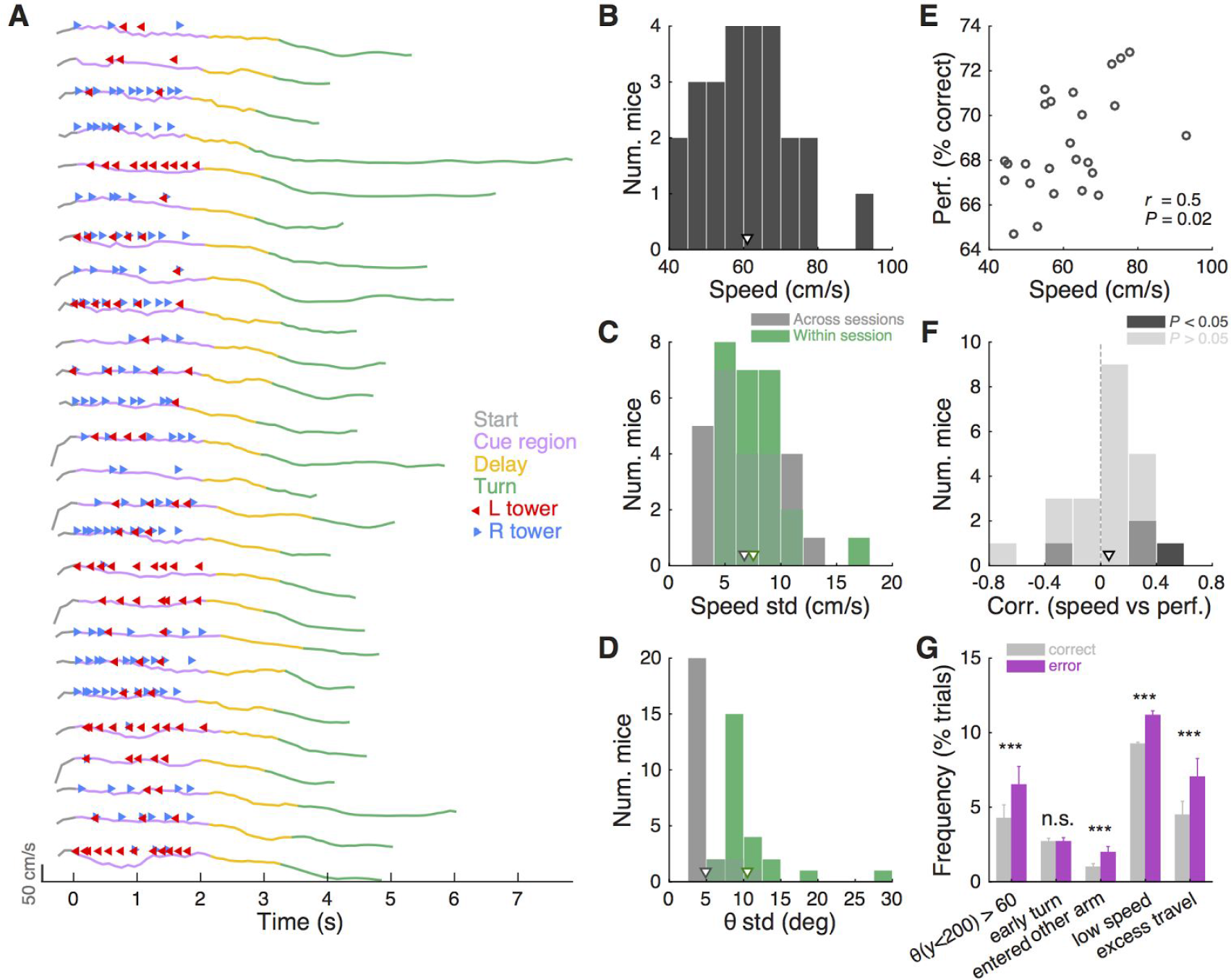
Stability of running patterns. **(A)** Examples of running speed over time during 25 consecutive trials, aligned by entry in the cue region (t = 0). Each line is color-coded according according to the portion of the maze (start, cue, delay or turn). Tower onset times are shown as leftward red or rightward blue arrows on top of each trace. **(B)** Distribution of average running speed across trials and sessions for animals with at least 1000 trials (n = 25). Arrowhead indicates population mean. **(C)** Distribution of standard deviations of average running speed across session-wide averages (gray, mean ± SEM: 6.7 ± 0.6 cm/s) and across trials within a session (green, mean ± SEM: 7.5 ± 0.6 cm/s). Arrowheads indicate population mean, and follow the same color code. **(D)** Distribution of standard deviations of average view angle across sessions and across trials within a session, calculated separately for right- and left-choice trials and then averaged (mean ± SEM: 4.9 ± 0.4 ˚ vs. 10.4 ± 0.9 ˚, respectively). Conventions as in C. **(E)** Correlation between average running speed and average overall performance across all sessions for each mouse (n = 25, *r* = 0.48, *P* = 0.02, Pearson's correlation). **(F)** Distribution of session-wise correlations between average running speed and average overall performance, showing that although there is an overall correlation between the two indicators, for any given mouse there is little correlation of speed and performance on individual sessions: only 4/25 mice had significant correlations between running speed and performance across sessions, and the sign of the correlation was negative for one of these mice (*r* = 0.06 ± 0.05, mean ± SEM). **(G)** Average frequency of different types of putative motor errors, belonging to five categories: trials with large-magnitude view angles during the cue period (> 60˚), trials with early turns (i.e. a turn immediately before the arm, resulting in a wall collision), trials in which the mouse first entered the opposite arm to its final choice, trials with speeds below the 10th percentile (defined separately for each mouse), and trials with traveled distance in excess of 110% of nominal maze length. Frequency was calculated separately for correct and error trials. Error bars, ± SEM. *** *P* < 0.001, n.s.: not significant.

**Supplementary Figure 8.**
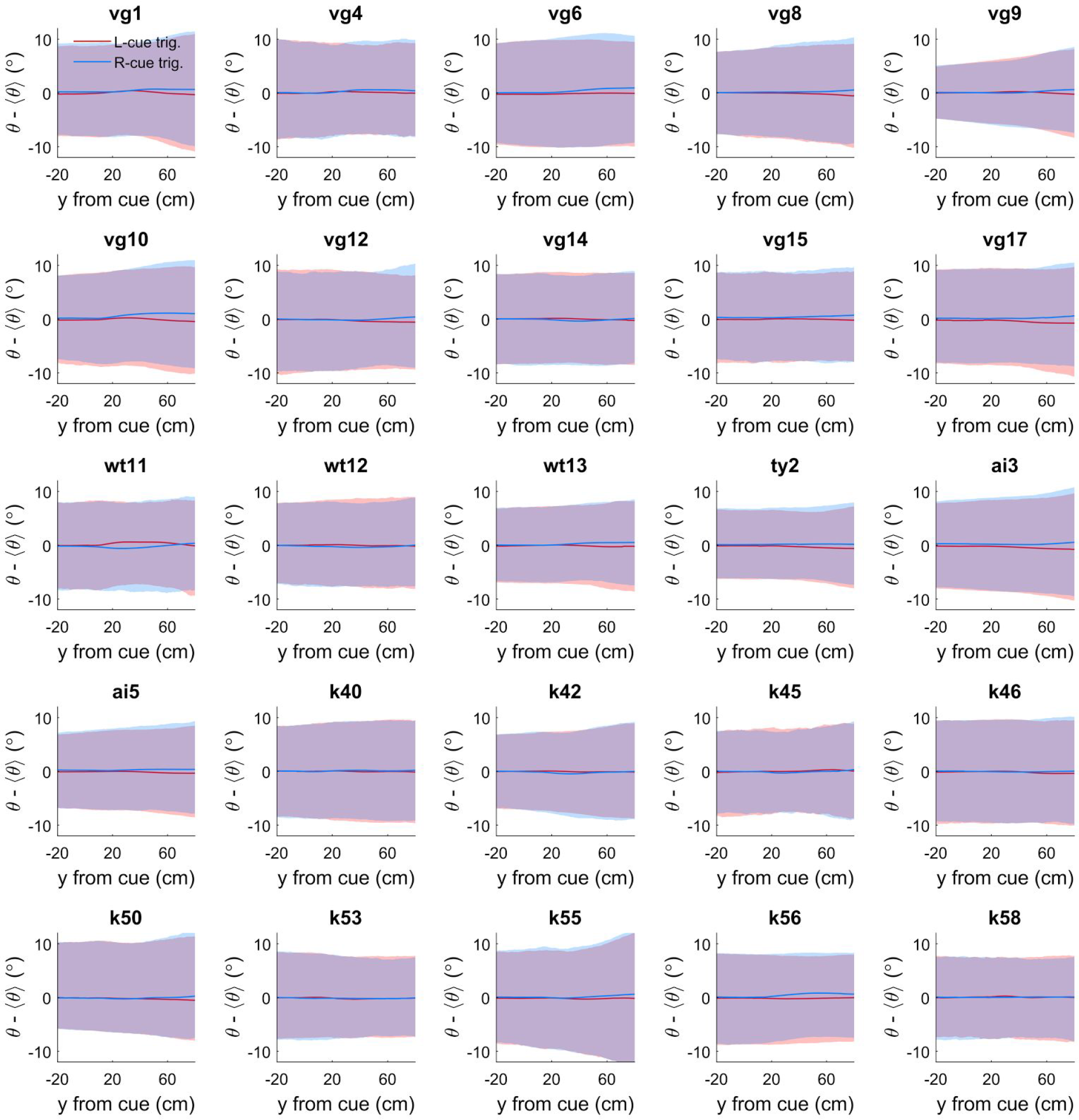
Cue-triggered change in view angles for individual mice. Each panel corresponds to data from a single mouse in this study, otherwise this is the same as Figure 8D: cue-triggered change in the view angle θ relative to the average trajectory 〈θ〉 for trials of the same choice. The bands indicate the 1 standard deviation spread across trials, with the lines being the mean across trials.

**Supplementary Figure 9.**
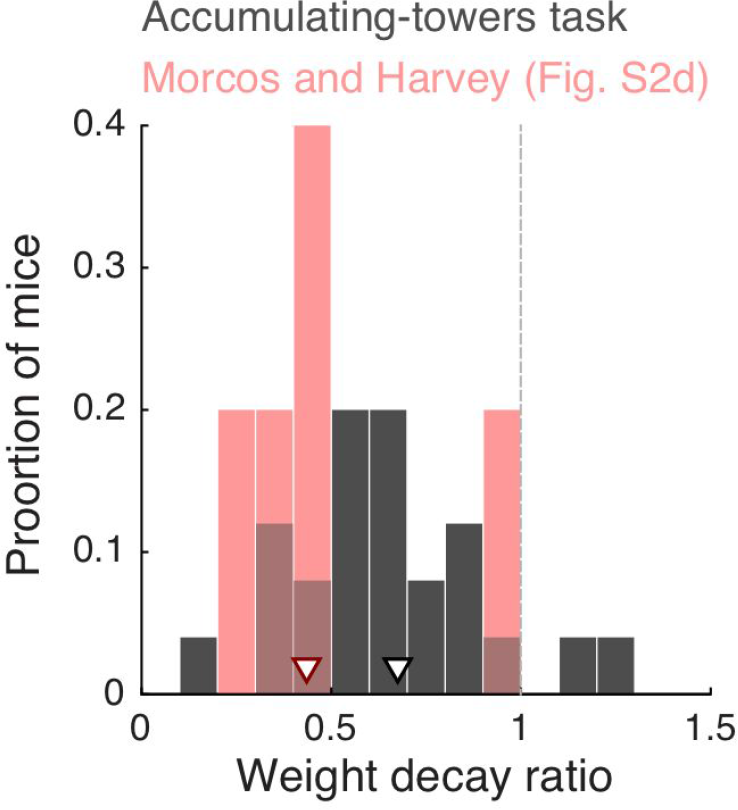
Comparison between the degree of primacy in the accumulating-towers and the Morcos and Harvey tasks. For direct comparison with the Morcos and Harvey task, we recalculated the logistic regression from the final accumulation maze of our task using 6 bins. Data from Supplementary Figure 2, panel d, in Morcos and Harvey (2016) was kindly provided by A.S. Morcos and C.D. Harvey. We then calculated the weight decay ratio as previously described (**Materials and Methods** and **Results, Fig. 3C**). Arrowheads, median.

**Supplementary Table 1.** Detailed parameters for all shaping mazes and main accumulation maze.

|  | T1 | T2 | T3 | T4 | T5 | T6 | T7 | T8 | T9 | T10 | T11 |
| --- | --- | --- | --- | --- | --- | --- | --- | --- | --- | --- | --- |
| **Total length (cm)** | 60 | 170 | 270 | 330 | 330 | 330 | 330 | 330 | 330 | 330 | 330 |
| **Cue period (cm)** | 45 | 120 | 220 | 280 | 280 | 240 | 200 | 200 | 200 | 200 | 200 |
| **Delay (cm)** | 10 | 20 | 20 | 20 | 20 | 60 | 100 | 100 | 100 | 100 | 100 |
| **Tower density (m<sup>-1</sup>)** | 3.0 | 3.8 | 3.8 | 3.8 | 4.2 | 4.2 | 4.2 | 4.5 | 4.8 | 4.8 | 5.0 |
| **Tower duration (ms)** | Inf | Inf | Inf | Inf | Inf | Inf | Inf | Inf | Inf | 200 | 200 |
| **Tower visible from (cm)** | 10 | 10 | 10 | 10 | 10 | 10 | 10 | 10 | 10 | 10 | 10 |
| **Visual guide?** | Y | Y | Y | Y | N | N | N | N | N | N | N |
| **Tower side ratio (m<sup>-1</sup>)** | Inf | Inf | Inf | Inf | Inf | Inf | Inf | 8.3:0.7 | 8.0:1.6 | 8.0:1.6 | 7.7:2.3 |
| **Warm-up** | none | none | none | none | 30 T4 trials with < 10% bias and > 80% correct | 30 T4 trials with < 10% bias and > 80% correct | 30 T4 trials with < 10% bias and > 80% correct | 10 T4 and 15 T7 trials with < 10% bias and > 80% correct | 10 T4 trials with < 10% bias and > 85% correct | 10 T4 trials with < 10% bias and > 85% correct | 10 T4 trials with < 10% bias and > 85% correct |
| **Advancement criteria** | 10 completed trials | 40 completed trials | 80 completed trials at > 60% correct | 2 sessions with 100 trials at > 90% correct | 1 session with 100 trials at > 80% correct | 1 session with 100 trials at > 80% correct | 1 session with 100 trials at > 80% correct | 1 session with 100 trials at > 75% correct | 1 session with 100 trials at > 70% correct | 1 session with 100 trials at > 70% correct | n/a |
| **Easy blocks**<br>(performance calculated over 40-trial window) | none | none | none | none | 10 T4 trials if < 70% correct | 10 T4 trials if < 70% correct | 10 T4 trials if < 70% correct | 10 T4 trials if < 65% correct | 10 T7 trials if < 60% correct | 10 T7 trials if < 60% correct | 10 T7 trials if < 55% correct |

**Supplementary Movie 1**. **Playback of six example trials from the accumulating-towers task. Left:** flattened view of the mice's perspective as they navigated the maze. The red lines indicate the estimated boundaries of the binocular field (± 17.5˚ at the horizon), and the yellow lines indicate ± 45˚ for reference. *θ*: view angle. Negative numbers indicate left side by convention. Luminance has been increased for convenience. **Right:** equivalent top-down view of the virtual maze. The mouse avatar turns according to its recorded virtual view angle, and towers become gray outlines when they disappear from the maze. Movie has been slowed down by 2x.

## Supplementary Methods

### Spatial Poisson distribution of tower locations

We used the following algorithm to randomly generate tower placement locations according to a Poisson process, i.e. with exponentially distributed inter-tower spacings subject to a minimum interval between towers:

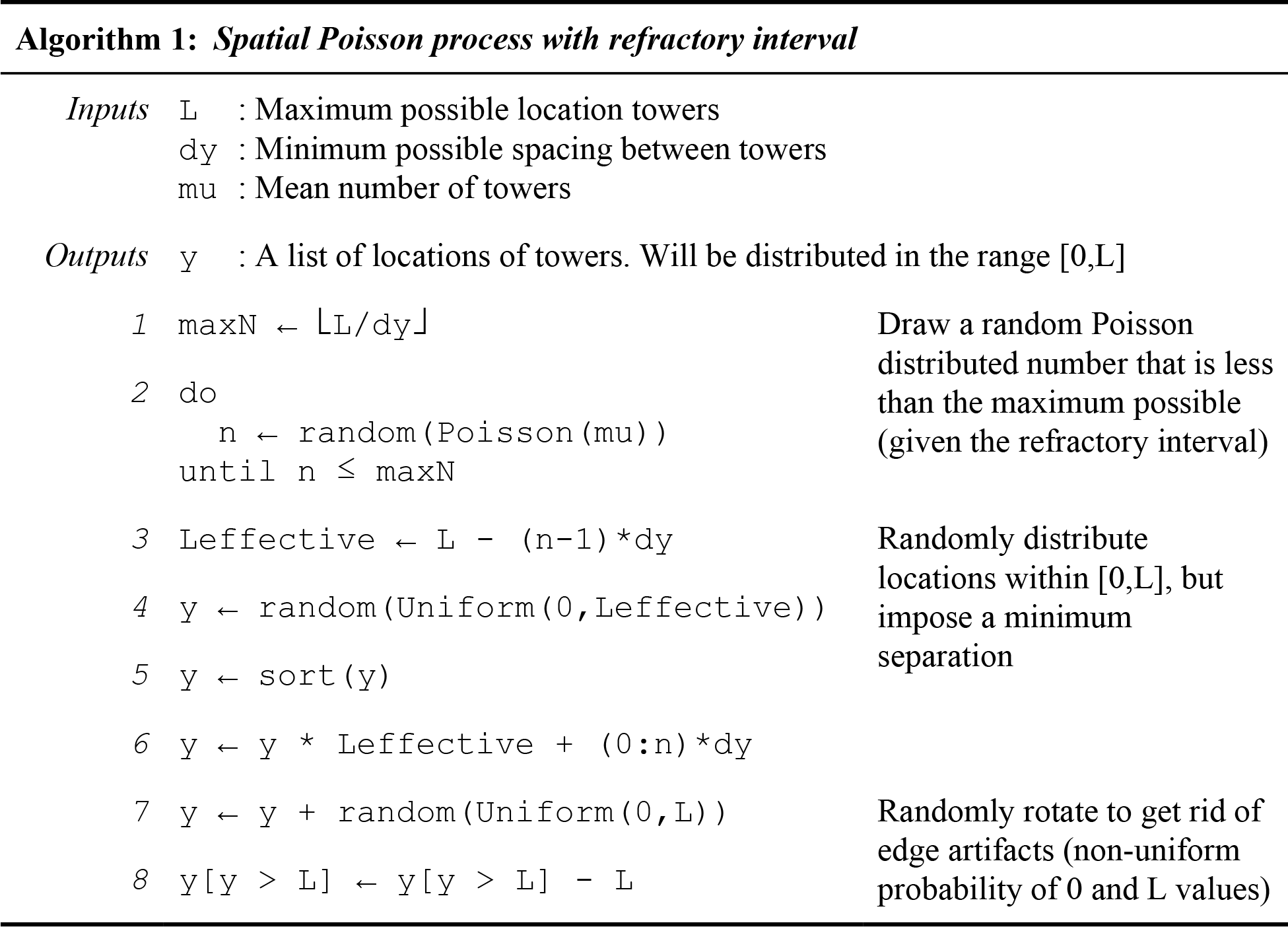

### Exponential gain for translating treadmill movements into changes in virtual view angle

Learning to use a spherical treadmill to execute navigational movements in virtual reality constitutes a substantial portion of the training time for this task. One of the optimizations we have performed to ease this process is to select treadmill-to-virtual-movement transformations so that mice can execute smooth motions without spending aversive amounts of time during turns into the arms of the T-maze. Historically we had first utilized a constant gain (Harvey et al. 2012) for the, but when this gain was low mice required a large amount of time to turn into the arms, encouraging them to initiate turns early (at the expense of accumulating later cues). Conversely, when this gain was high, small postural shifts in the stem of the T-maze caused the virtual scene to wobble, which was undesirable in a task involving visual cues. These observations motivated the use of a nonlinear gain function that deemphasizes small, uncontrollable movements of the treadmill during running down the stem, but facilitates sharper turns at the end of the T-maze to encourage straighter view angle trajectories.

### Heuristic models: optimization technique

Here we defined several models where the choice of the mouse in a series of trials is assumed to be a Bernoulli process parameterized by a probability of making a choice to the right, 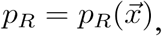 that depends on a set of trial-specific quantities 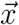 (see **Materials and Methods**).

We obtained best-fit parameters for each model by maximizing the log likelihood of the model for a given dataset comprising *m* of trials. Let the mouse’s choice on the i^th^ trial be *c_i_, i* = 1, …, *m* which is 1 (0) if the mouse chose right (left), then the likelihood of observing this choice is given by the binomial distribution 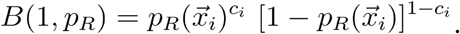 Taking the product of individual-trial likelihoods we obtain:

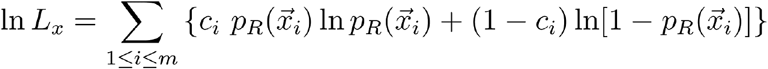

Additionally we subtracted L1 penalty terms for all free parameters of the model. For a model that includes all factors, the quantity that is maximized is therefore:

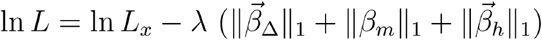

where 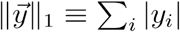 is the L1 norm. This regularization is used as a method for selecting the most parsimonious model in terms of driving coefficients to zero when they do not result in a significantly better fit for the model (Schmidt, 2010). It was also crucial for some models, particularly those that contain history-dependent lapse terms, because of the presence of multiple local maxima that made the problem otherwise ill-posed.

The regularization strength hyperparameter λ was determined by using a 3-fold cross-validation (CV) procedure to find the optimal model in terms of predictive power. A given dataset was first divided into thirds, and each third is used exactly once as a test set and the remaining two thirds as its complementary training set. To equalize the highly different scales of the 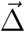 factors compared to the rest of the factors which are bounded within [-1,1], for each coordinate the standard deviation 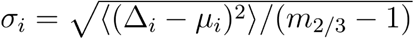 was computed using the trials in the training set, and used to scale the evidence factors, Δ_i_ → Δ_i_/σ_i_. In other words, the only thing that this changed was that the coefficients 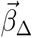 were expressed in units of 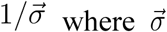 are constants derived using the training set (the same are used for the test set, as it would be unfair use of information if they were re-derived for the test set).

### Alternative strategy models: one-random-tower analysis details

For the analysis in **Figure 4C**, for each mouse we selected the top 1 third performance blocks, and only analyzed mice that had at last 200 trials in these blocks; we pooled together all trials from these blocks and mice. To test the 1-random tower hypothesis, we reasoned that we expect to obtain a linear psychometric curve when the sum of towers (#R+#L) was fixed for all trials. This is because the probability to go right for the 1-random tower strategy is given by #R/(#R+#L), and if the denominator is fixed, then the psychometric curve (which is given by (#R-#L)/(#R+#L)) is linear in the difference of towers #R-#L, which is the standard x-axis of the psychometric curve. However, we have empirically observed sigmoid shapes for the psychometric curves of the mice's choices. Thus, we proceeded to quantify if the psychometric curves of the mice choices were different from that of the 1-random tower model (as described in **Materials and Methods**). To obtain a dataset with fixed #R+#L, we next selected only trials where #R+#L=12. This number was chosen because it was the maximum number of #R+#L for which there were at least 4000 trials. We then found the psychometric curve for the actual data and the 1-random tower model. As expected, the 1-random tower model results in a linear psychometric curve, whereas the actual data appears more sigmoidal. To find whether these curves are significantly different from each other, we performed a shuffling test in the following way: we generated 5000 bootstrapped pairs of curves by pooling for all trials with a given #R – #L the number of times the mice (or model) chose right, and then randomly assigning the same number of right choices between the two curves, while keeping the total number of trials as in the original data. The sum of absolute differences between the two curves was used as the test statistic.

